# Reduced expression of C/EBPβ-LIP extends health- and lifespan in mice

**DOI:** 10.1101/250100

**Authors:** Christine Müller, Laura M. Zidek, Tobias Ackermann, Tristan de Jong, Peng Liu, Verena Kliche, Mohamad Amr Zaini, Gertrud Kortman, Liesbeth Harkema, Dineke S. Verbeek, Jan P. Tuckermann, Julia von Maltzahn, Alain de Bruin, Victor Guryev, Zhao-Qi Wang, Cornelis F. Calkhoven

## Abstract

Ageing is associated with physical decline and the development of age-related diseases such as metabolic disorders and cancer. Few conditions are known that attenuate the adverse effects of ageing, including calorie restriction (CR) and reduced signalling through the mechanistic target of rapamycin complex 1 (mTORC1) pathway. Synthesis of the metabolic transcription factor C/EBP β ‐LIP is stimulated by mTORC1, which critically depends on a short upstream open reading frame (uORF) in the C/EBP β-mRNA. Here we describe that reduced C/EBP β ‐LIP expression due to genetic ablation of the uORF delays the development of age-associated phenotypes in mice. Moreover, female C/EBP β^ΔuORF^ mice display an extended lifespan. Since LIP levels increase upon aging in wt mice, our data reveal an important role for C/EBPβ in the aging process and suggest that restriction of LIP expression sustains health and fitness. Thus, therapeutic strategies targeting C/EBP β ‐LIP may offer new possibilities to treat age-related diseases and to prolong healthspan.

## Introduction

Delaying the occurrence of age related-diseases and frailty (disabilities) and thus prolonging healthspan, would substantially increase the quality of life of the ageing population and could help to reduce healthcare costs. Calorie restriction (CR) or pharmacological inhibition of the mTORC1 pathway by rapamycin are considered as potential effective interventions to delay aging and to increase healthspan in different species (Kaeberlein, Rabinovitch, & Martin, 2015). However, for humans CR is a difficult practice to maintain and may have pleiotropic effects depending on genetic constitution, environmental factors and stage of life. Likewise, the long-term use of rapamycin is limited by the risk of side effects, including disturbed glucose homeostasis, impaired would healing, gastrointestinal discomfort and others (Augustine, Bodziak, & Hricik, 2007; de Oliveira et al., 2011; Lamming et al., 2012; Wilkinson et al., 2012). Therefore, there is a need to investigate alternative targets that are part of the CR/mTORC1 pathway that can be manipulated to reach similar beneficial effects. Our work suggests that the transcription factor C/EBPβ may provide such a target.

C/EBPβ regulates the expression of metabolic genes in liver and adipose tissue (Desvergne, Michalik, & Wahli, 2006; Roesler, 2001). From its mRNA three protein isoforms are synthesized through the usage of different translation initiation sites: two isoforms acting as transcriptional activators, Liver Activator Protein (LAP) -1 and -2, and a transcriptional inhibitory isoform called Liver-enriched Inhibitory Protein (LIP) (Descombes & Schibler, 1991). We showed earlier that translation into LIP depends on a *cis*-regulatory uORF (**Figure 1A**) and is stimulated by mTORC1 signalling (Calkhoven, Muller, & Leutz, 2000; Jundt et al., 2005; Zidek et al., 2015). Pharmacological or CR-induced inhibition of mTORC1 in mice selectively reduces LIP-protein synthesis and thereby increases the LAP/LIP ratio in different tissues, which is associated with metabolic improvements resembling CR (Albert & Hall, 2015; Zidek et al., 2015). Recently we have developed a screening strategy for the discovery of small molecule drugs that suppress LIP expression, demonstrating that LIP expression is therapeutically targetable (Zaini et al., 2017).

**Figure 1.**
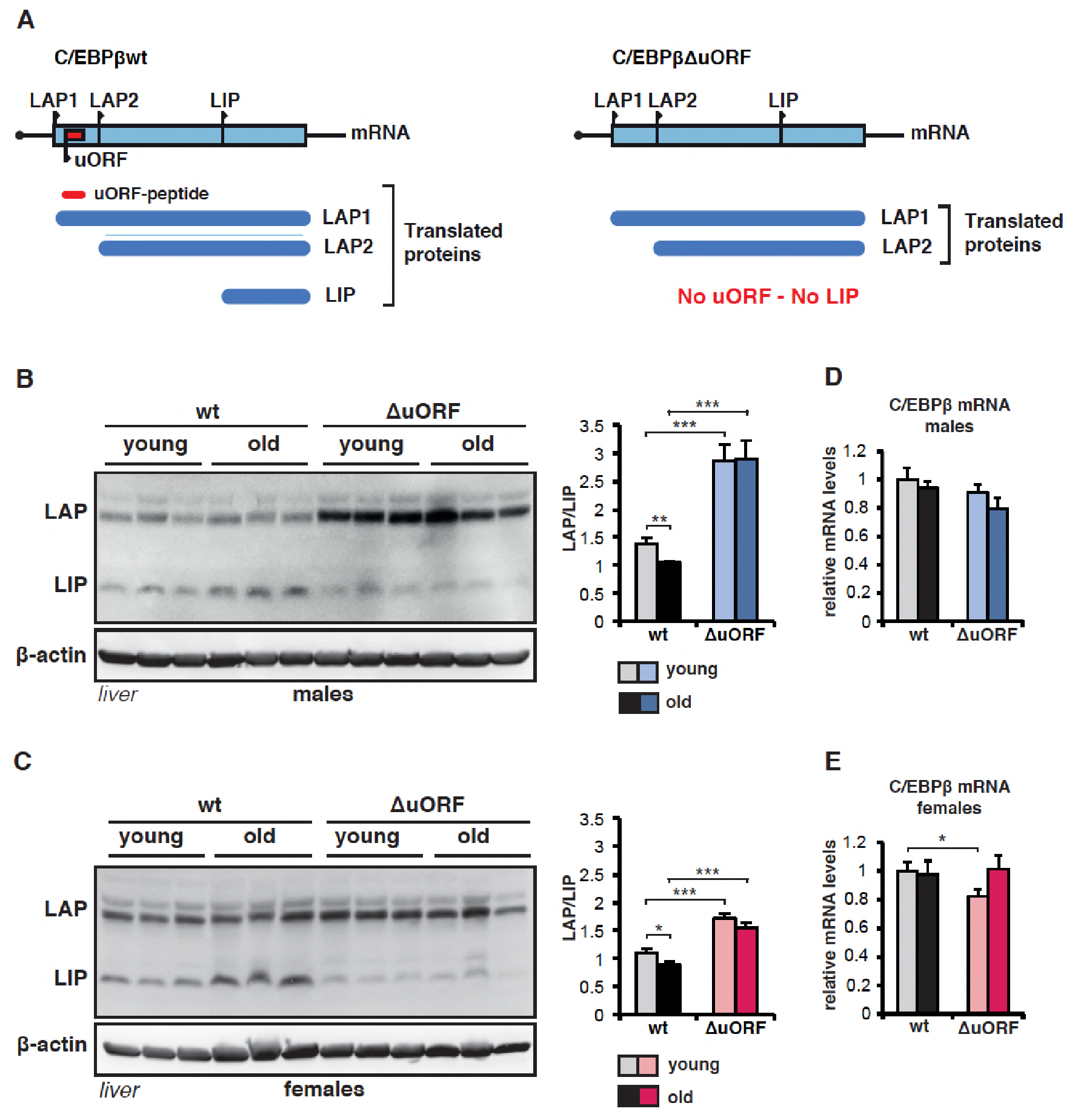
C/EBPβ LAP/LIP isoform ratio increases upon ageing. (A) The graph at the left shows that wt C/EBPβ-mRNA is translated into LAP1 and LAP2 through regular translation initiation, while translation into LIP involves a primary translation of the uORF followed by translation re-initiation at the downstream LIP-AUG by post-uORF-translation ribosomes. The graph at the right shows that genetic ablation of the uORF abolishes translation into LIP, but leaves translation into LAP1 and LAP2 unaffected (for detailed description see(Calkhoven et al., 2000; Zidek et al., 2015). (B and C) Immunoblots of liver samples from young (5 months) and old (female 20 months, male 22 months) wt and C/EBPβ^ΔuORF^ (B) males and (C) females showing LAP and LIP isoform expression. β-actin expression served as loading control. The LAP/LIP isoform ratio as calculated from quantification by chemiluminescence digital imaging of immunoblots is shown at the right (wt males n=9 young, n=10 old; C/EBPβ^ΔuORF^ males, n=11 young, n=10 old; wt females, n=9 wt young, n=8 old; C/EBPβ^ΔuORF^ females, n=10 young, n=10 old). (D) C/EBPβ mRNA levels in males as determined by quantitative real-time PCR (wt, n=11 young, n=11 old; C/EBPβ^ΔuORF^, n=11 young, n=9 old) and (E) in females (wt, n=9 young, n=11 old; C/EBPβ^ΔuORF^, n=9 young, n=11 old). P-values were determined by Student’s t-test, *p<0.05; **p<0.01; ***p<0.001.

## Results and discussion

Others showed that LIP levels increase during aging in liver and white adipose tissue (WAT) (Hsieh et al., 1998; Karagiannides et al., 2001; Timchenko et al., 2006). Similarly, in our cohorts of wt C57BL/6 mice LIP levels are significantly higher in livers of old (20-22 months) versus young (5 months) mice, resulting in a decrease in LAP/LIP ratios during ageing (**Figure 1B, C - figure supplement 1**). In contrast, in C/EBPβ^ΔuORF^ mice LIP levels are low and stay low in old mice. LAP levels in C/EBPβ^ΔuORF^ males and to a lesser extend in females are increased, which is probably due to additional initiation events at the LAP-AUG by ribosomes that normally would have initiated at the uORF (Calkhoven et al., 2000). The C/EBPβ-mRNA levels are comparable at different ages and in the different genotypes (**Figure 1D and E**). Thus, LIP levels increase with age and this increase is dependent on the uORF in the C/EBPβ-mRNA.

We hypothesised that the C/EBPβ^ΔuORF^ mutation may have positive effects on healthspan and lifespan based on the CR-like metabolic improvements in C/EBPβ^ΔuORF^ mice, including enhanced fatty acid oxidation and lack of steatosis, improved insulin sensitivity and glucose tolerance and higher adiponectin levels (Zidek et al., 2015). A lifespan experiment was set up comparing C/EBPβ^ΔuORF^ mice with wt littermates (C57BL/6) in cohorts of 50 mice of each genotype and gender. The survival curves revealed an increase in median survival of 20.6% (difference in overall survival p=0.0014 log-rank test, n=50) for the female C/EBPβ^ΔuORF^ mice compared to wt littermates (**Figure 2A**). From the 10% longest-lived females, nine out of ten were C/EBPβ^ΔuORF^ mice (**Table supplement 1**), showing that the maximum lifespan of C/EBPβ^ΔuORF^ females is significantly increased (p=0.0157 Fisher’s exact test). If maximum lifespan is determined by the mean survival of the longest-lived 10% of each cohort, C/EBPβ^ΔuORF^ females show an increase of 9.14% (p-value=0.00105 Student’s ttest.). For the male cohort we observed a modest increase in median survival of 5.2%, however the overall survival was not significantly increased (p=0.4647 log-rank test, n=50) (**Figure 2B**). The increase in median survival of the combined cohort of C/EBPβ^ΔuORF^ mice (males & females) was 10.5% (with a significant increase in overall survival p=0.0323 log-rank test, n=100) (**figure supplement 2 and Table supplement 1**). The observed median survival for wt females (623 days) is lower than what most other labs have reported for C57BL/6 females. We reasoned that this was due to a high incidence of ulcerative dermatitis (UD) we observed particularly in our female cohort (females: 19 mice or 38% for wt and 26 mice or 52% for C/EBPβ^ΔuORF^; males: 15 mice or 30% for wt and 10 mice or 20% for C/EBPβ^ΔuORF^). UD is a common and spontaneous condition in mice with a C57BL/6 background that progress to a severity that euthanasia is inevitable (Hampton et al., 2012). Therefore, survival curves were also calculated separately for UD-free mice and for mice that were euthanized because of serious UD (**Figure 2 C-F** and **Table supplement 1** for complete overview). These data show that median lifespan of UD-free wt females are in a more normal range (740 days) and that the C/EBPβ^ΔuORF^ mutation results in a significant increase of median survival specifically in females irrespective of the condition of UD. Moreover, the median survival of the C/EBPβ^ΔuORF^ UD-free females (860.5 days) is higher compared to both wt females and wt males (829 days). The separate analysis of males with or without UD in both cases didn’t show any significant difference in lifespan (**Figure 2D and F** - **figure supplement 2B and C**) similarly to the analysis of the whole cohort indicating that the C/EBPβ^ΔuORF^ mutation increases lifespan specifically in females.

**Figure 2.**
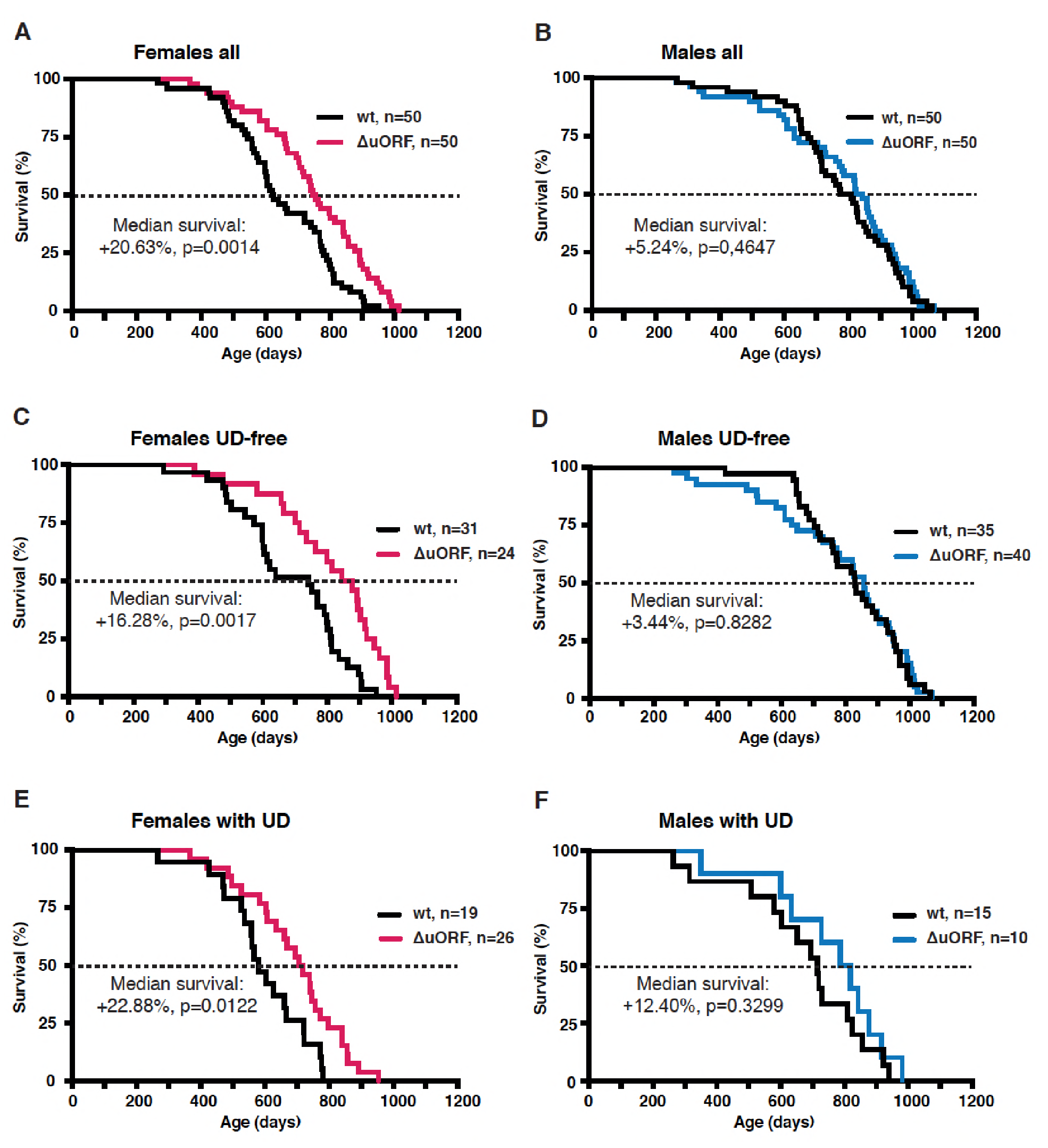
Increased survival of female C/EBPβ^ΔuORF^ mice. Survival curves of (A) the complete female cohorts, (B) complete male cohorts, (C) the UD-free female cohorts, (D) UD-free male cohorts, (E) female mice with UD and (f) male mice with UD with the survival curves of wt or C/EBPβ^ΔuORF^ mice indicated. The increase in median survival (%) of C/EBPβ^ΔuORF^ compared to wt littermates and statistical significance of the increase in the overall survival as determined by the log-rank test is indicated in the figure.

Similar to our study several other studies report on sex specific differences in lifespan regulation. Calorie restriction by 20% has a greater lifespan extending effect in female C57BL/6 or DBA/2J mice compared to males (Mitchell et al., 2016). In addition, prolonged treatment with the mTORC1-inhibitor rapamycin starting from midlife has lifespan extending effects that are much stronger in females than in males using genetically heterogeneous mice as well as C57BL/6 mice (Fok et al., 2014; Harrison et al., 2009; Miller et al., 2011; Y. Zhang et al., 2014). On the contrary, transient rapamycin treatment for 90 days during midlife induces extension of lifespan in males but not in females depending on the dose (Bitto et al., 2016). Moderate overexpression of the mTORC1-inhibitor TSC1 or deletion of the ribosomal S6 protein kinase 1 (S6K1), downstream of mTORC1, result in lifespan extension only in females (Selman et al., 2009; H. M. Zhang, Diaz, Walsh, & Zhang, 2017). Mutations affecting both mTORC1 and mTORC2 show ambiguous effects; lifespan extension was limited to females in mice heterozygous for mTOR and its cofactor mammalian lethal with Sec 13 protein 8 (mLST8) (Lamming et al., 2012), while in a mTOR-hypomorphic mouse model lifespan is extended in both males and females (Wu et al., 2013). Notably, the reduction of LIP expression under low mTORC1 signalling is dependent on 4E-BP1/2 function and not on inhibition of S6K1 (Zidek et al., 2015). Thus, the bias towards female lifespan extension upon reduced mTORC1 signalling seems to be a common feature irrespective of whether the S6K1 or 4E-BP branch is affected. Also in mouse strains with alterations in other pathways like the somatotropic axis lifespan extension is often, but not always, more pronounced in females (Brown-Borg, 2009). Examples of somatotropic-related female biased lifespan extension are Ames dwarf mice that are deficient in growth hormone (GH) and prolactin production (Brown-Borg, Borg, Meliska, & Bartke, 1996) and insulin-like growth factor 1 (IGF-1) receptor heterozygous mice The reason for the mostly female biased lifespan extension upon inhibition of mTORC1 signalling or in the other mentioned mouse models is not known.

Aging is the most important risk factor for development of cancer. A reduction in cancer incidence is recurrently observed upon CR, rapamycin-treatment or manipulation of other pathways that increase longevity in several animal models (Anisimov et al., 2011; Colman et al., 2009; Komarova et al., 2012; Mattison et al., 2012; Neff et al., 2013; Serrano, 2016; Weindruch & Walford, 1982). Mice in the lifespan cohorts that died or were sacrificed according to humane endpoint criteria underwent necropsy and tumours were analysed by a board certified veterinary pathologists of the Dutch Molecular Pathology Centre (DMPC). The incidence of neoplasms was markedly reduced in female C/EBPβ^ΔuORF^ mice compared to female wt mice (68% -> 45,8%, p=0.025 Fisher’s exact test) (**Figure 3A**). Furthermore, tumours were detected on necropsy at a higher age in female C/EBPβ^ΔuORF^ mice compared to wt mice indicating a delay in tumour development (**Figure 3B**). The increase in median survival of the tumour bearing C/EBPβ^ΔuORF^ females was 25.49% compared to that of tumour bearing wt females (p=0.0217 log-rank test) (**figure supplement 3A**). Also the tumour load (number of different tumour types per mouse) and the tumour spread (total number of differently located tumours per mouse irrespective of the tumour type) were lower in female C/EBPβ^ΔuORF^ mice (**figure supplement 3B**). For males no significant reduction in tumour incidence was detected in C/EBPβ^ΔuORF^ mice (**Figure 3C and 3D**). The survival of tumour bearing mice and the tumour load was similar in wt and C/EBPβ^ΔuORF^ males, while the tumour spread seems to be even slightly increased in C/EBPβ^ΔuORF^ male mice (**figure supplement 3C, D**). The main tumour types found in female mice were lymphoma, hepatocellular carcinoma and histiocytic sarcoma. The occurrence of all three types was reduced in C/EBPβ^ΔuORF^ females (**Table supplement 2**). For other tumour types the single numbers are too small to make a clear statement about a change in frequency. In male mice hepatocellular carcinoma and histiocytic sarcoma were the most frequent tumour types observed. Although the overall tumour incidence was similar in C/EBPβ^ΔuORF^ and wt males, the frequency of hepatocellular carcinoma was reduced in the C/EBPβ^ΔuORF^ males (**Table supplement 2**). Notably, knockin mice with elevated LIP levels show an increased tumour incidence upon ageing that goes along with reduced survival compared to wt controls (Begay et al., 2015). LIP overexpression can stimulate tumour cell proliferation, migration and transformation in vitro and high LIP levels have been detected in different human tumour tissues (Anand et al., 2014; Arnal-Estape et al., 2010; Calkhoven et al., 2000; Haas et al., 2010; Jundt et al., 2005; Park et al., 2013; Raught et al., 1996; Zahnow, Younes, Laucirica, & Rosen, 1997). Taken together the studies support an oncogenic role of LIP in tumour development and suggest that the reduction of LIP in the C/EBPβ^ΔuORF^ mice counteracts tumour development at least partially by cell intrinsic mechanisms.

**Figure 3.**
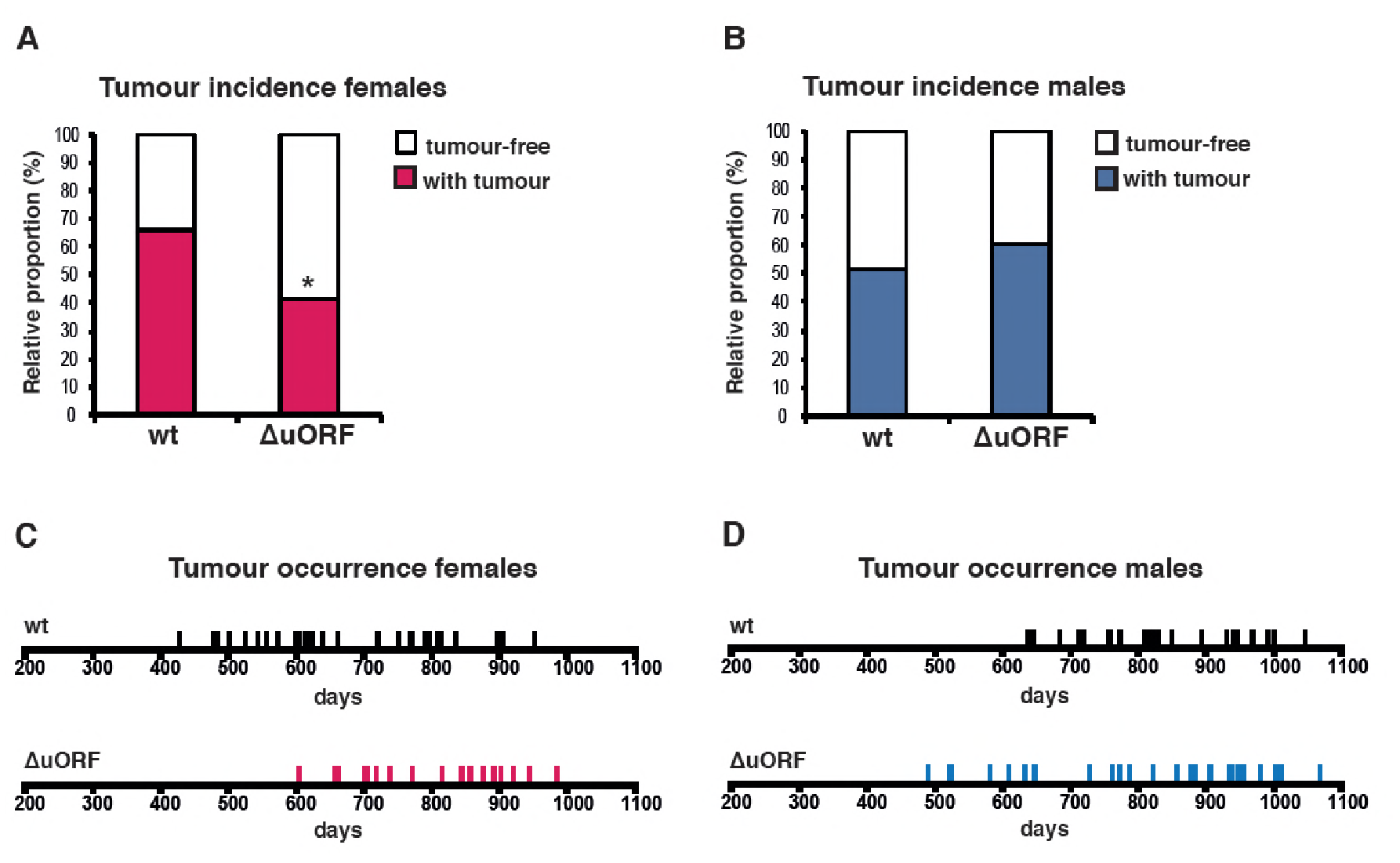
Reduced incidence and delayed occurrence of tumours is female C/EBPβ^ΔuORF^ mice. (A) Tumour incidence of females as determined by pathological examination of neoplasms found upon necropsy of mice from the lifespan cohorts (wt, n= 50; C/EBPβ^ΔuORF^, n=48). Statistical significance was calculated using Fisher’s exact test with *p<0.05. (B) Tumour incidence of males as determined by pathological examination of neoplasms found upon necropsy (wt, n=47; C/EBPβ^ΔuORF^, n=45. (C) Tumour occurrence in the female lifespan cohorts upon necropsy is shown for wt (black lines) and C/EBPβ^ΔuORF^ mice (red lines). (D) Tumour occurrence in the male lifespan cohorts upon necropsy is shown for wt (black lines) and C/EBPβ^ΔuORF^ mice (blue lines).

Apart from the reduced tumour incidence and the increase in survival of tumour-bearing C/EBPβ^ΔuORF^ females, also the survival of tumour-free female C/EBPβ^ΔuORF^ mice was significantly extended by 25.13% (p=0.0467 log-rank test) compared to wt tumour-free females (**figure supplement 3E**). This suggests that both the tumour incidence and additional unrelated factors contribute to the increased survival of C/EBPβ^ΔuORF^ females. The observed increase in median lifespan of tumour-free C/EBPβ^ΔuORF^ males of 19.71% does not correlate with a statistically significant increase in the overall survival (p=0.4647 log-rank test) (**figure supplement 3F**). However, the survival curve points to a possible health improvement in the median phase of the male lifespan. Taken together, the C/EBPβ^ΔuORF^ mutation in mice restricting the expression of LIP results in a significant lifespan extension and decreased tumour incidence in females but not in males.

Typically, CR-mediated, genetic or pharmacological suppression of mTORC1 signalling is accompanied by the attenuation of an age-associated decline of health parameters (Johnson, Rabinovitch, & Kaeberlein, 2013). We examined the selected health parameters of body weight and composition, glucose tolerance, naïve/memory T-cell ratio, motor coordination and muscle strength in separate ageing cohorts of young (3-5 months) and old (18-20 months for females and 20-22 months for males) mice. In addition, we compared the histological appearance of selected tissues (liver, muscle, pancreas, skin, spleen and bone) between old (20/22 months) wt and C/EBPβ^ΔuORF^ mice. Body weight was significantly increased in all old mice (**Figure 4A, B**). The increase for the old female C/EBPβ^ΔuORF^ mice was significantly smaller compared to old wt littermates, while for the males there was no significant difference between the genotypes (**Figure 4A, B**). The slightly lower body weight for the young C/EBPβ^ΔuORF^ males was also observed in our previous study (Zidek et al., 2015). A similar pattern was observed regarding the fat content that was measured by abdominal computed tomography (CT) analysis (**Figure 4C, D**). The volumes of total fat increased strongly in old mice both in visceral and subcutaneous fat depots (**figure supplement 4A, B**). Old female C/EBPβ^ΔuORF^ mice accumulated significantly less fat in the visceral and subcutaneous fat depots than wt females, while there was no difference for male mice (**figure supplement 4A, B**). The lean body mass was slightly lower in old female C/EBPβ^ΔuORF^ mice and increased in male wt mice compared to young mice (**figure supplement 4A, B**). Thus, female C/EBPβ^ΔuORF^ mice gain less fat upon aging similar to mice under CR or upon prolonged rapamycin treatment (Fang et al., 2013) (**figure supplement 4C**), which probably contributes to an improved health state upon aging and to the extension in lifespan. In contrast, although male C/EBPβ^ΔuORF^ mice had a lower body weight and subcutaneous fat content at a young age compared to wt mice they were not able to maintain this difference during the aging process, which correlates with the lack in lifespan extension. In addition, we found an increase in mRNA expression of the macrophage marker CD68 as a measure for age-related macrophage infiltration in visceral WAT of old mice, which was attenuated in female C/EBPβ^ΔuORF^ mice but not in male C/EBPβ^ΔuORF^ mice (**figure supplement 4D**).

**Figure 4.**
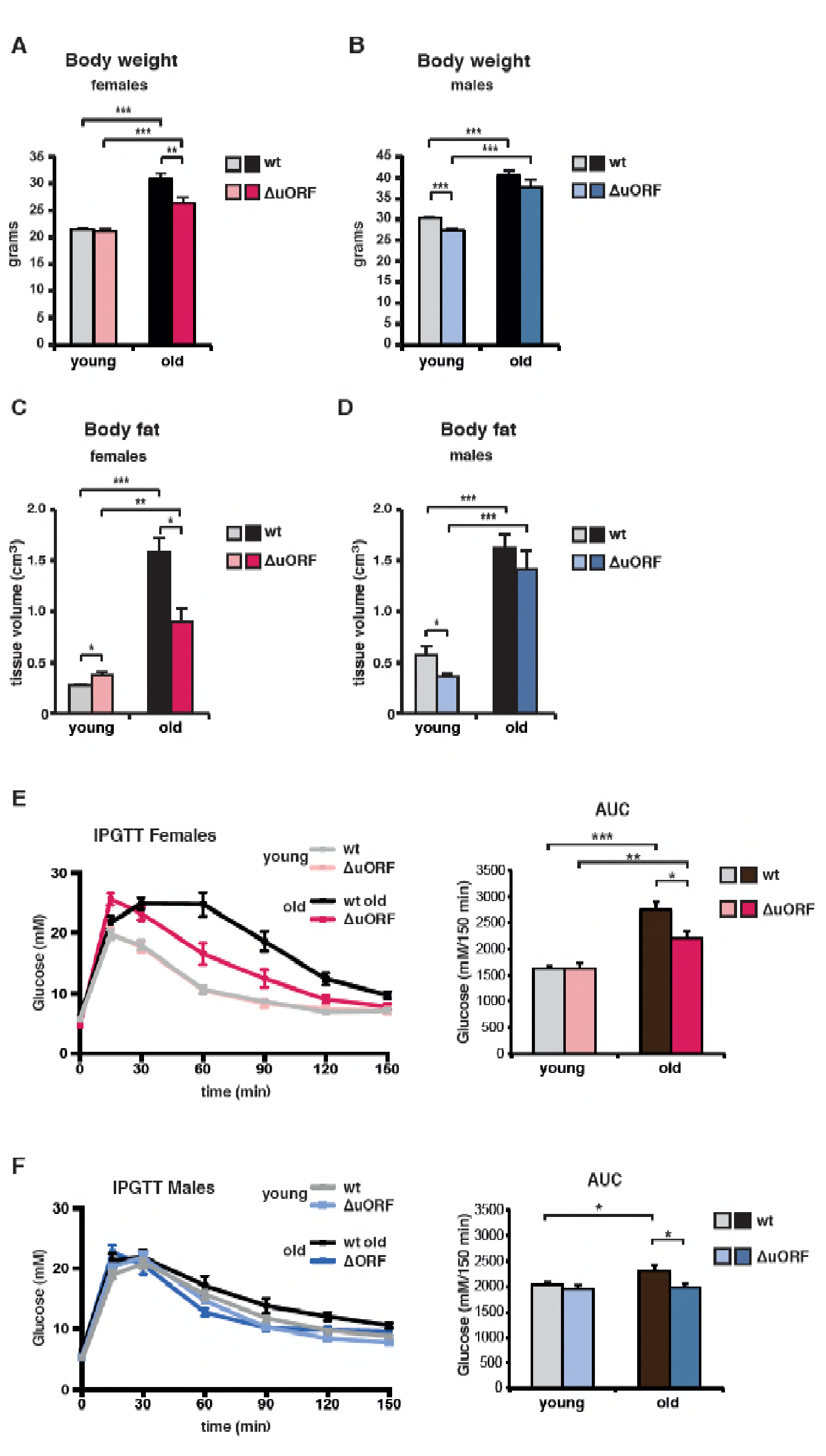
Ageing-associated increase in body weight, fat content and glucose tolerance is attenuated in female C/EBPβ^ΔuORF^ mice. (A) Body weight (g) of young (4 months) and old (19 months) female mice (wt, n=11 young, n=12 old; C/EBPβ^ΔuORF^, n=11 young, n=12 old). (B) Body weight of young (4 months) and old (21 months) male mice (wt, n=12 young and old; C/EBPβ^ΔuORF^, n=12 young, n=11 old). (C) Body fat content (cm^3^) as determined by CT analysis of young (4 months) and old (19 months) female mice (wt, n=11 young and old; C/EBPβ^ΔuORF^, n=9 young, n=11 old). (D) Body fat content of young (4 months) and old (21 months) male mice (wt, n=12 young and old wt; C/EBPβ^ΔuORF^, n=11 young, n=9 old). (E and F) i.p.-Glucose Tolerance Test (IPGTT) was performed with young (4 months) and old (female 19 months, male 21 months) wt and C/EBPβ^ΔuORF^ (E) females and (F) males. The area under the curve (AUC) at the right shows the quantification (wt females, n=9 young, n=10 old; C/EBPβ^ΔuORF^ females, n=10 young, n=11 old; wt males, n=12 young, n=11 old; C/EBPβ^ΔuORF^ males, n=11 young, n=10 old). P-values were determined by Student’s t-test, *p<0.05; **p<0.01; ***p<0.001.

Impaired glucose tolerance is a hallmark of the aging process, which is improved by CR (Barzilai, Banerjee, Hawkins, Chen, & Rossetti, 1998; Mitchell et al., 2016). The intraperitoneal glucose tolerance test (IPGTT) showed that glucose clearance, calculated as the area under the curve (AUC), is significantly less efficient in old wt compared to young wt mice (**Figure 4E, F**). Old C/EBPβ^ΔuORF^ females and males perform significantly better in the IPGTT test than old wt littermates, which is reflected by the lower AUC value. Therefore, the C/EBPβ^ΔuORF^ mutation protects against age-related decline of glucose homeostasis in males and females.

The ageing associated increase in memory/naïve T-cell ratio is a robust indicator for the progression of the immunological ageing progress. At a young age naïve T cells predominate and memory T cells are relatively scarce. Upon ageing the naïve T cell population is strongly reduced with a concomitant increase in the memory T cell population, resulting in an increased ratio of memory to naïve T cells (Hakim, Flomerfelt, Boyiadzis, & Gress, 2004). The ratio of memory (CD44high) to naïve (CD44low/CD62Lhigh) cytotoxic T (CD8+) cells or memory (CD44high) to naïve (CD44low/CD62Lhigh) helper T (CD4+) cells was analysed by flow cytometric analysis. Both increased upon aging in the blood of males and females of both genotypes (**Figure 5A-D**). However in C/EBPβ^ΔuORF^ mice of both genders this increase was significantly attenuated compared to wt mice (**Figure 5A-D - figure supplement 5A-D**). These data suggest that the C/EBPβ^ΔuORF^ mutation preserves a more juvenile immunological phenotype during ageing.

**Figure 5.**
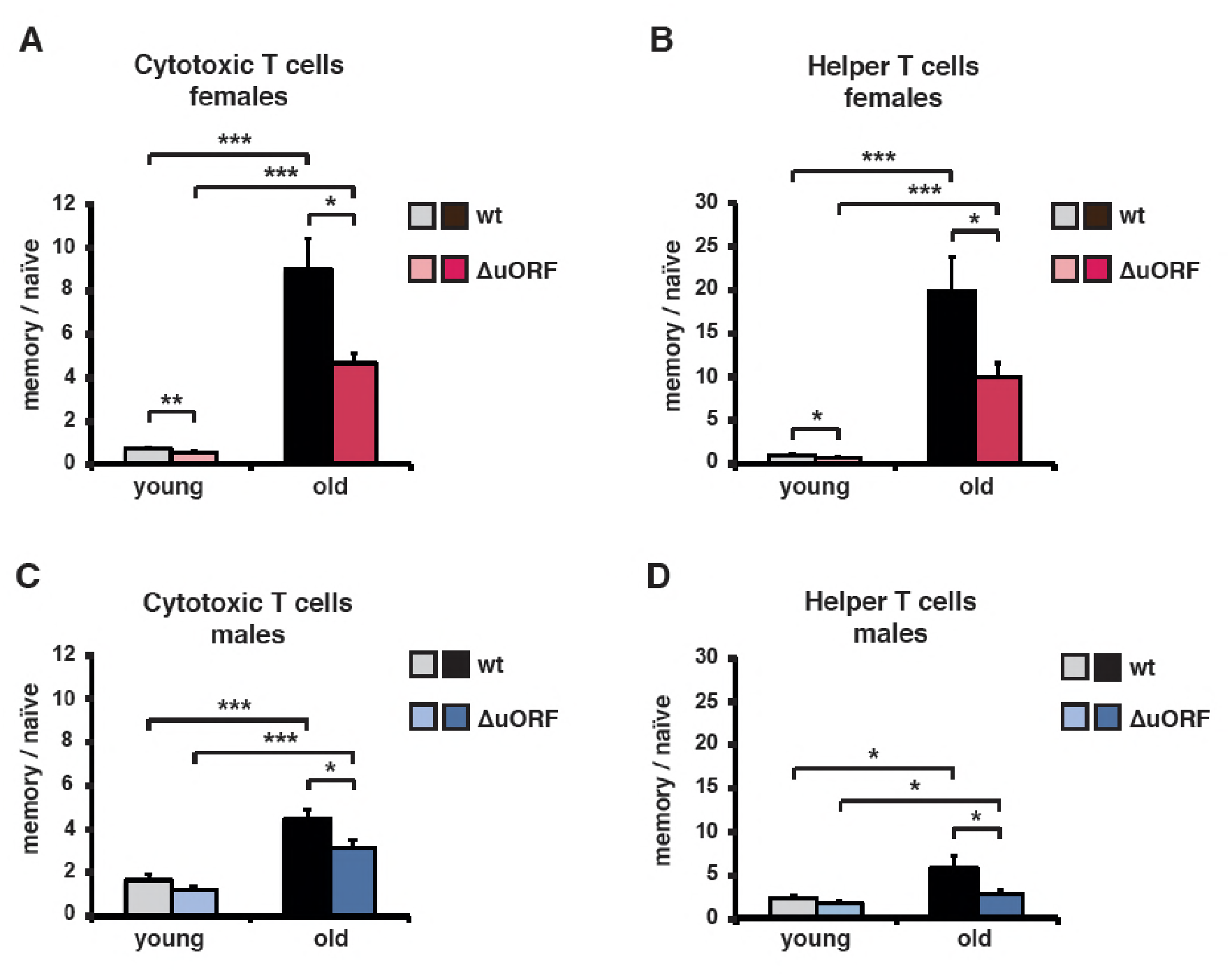
Ageing-associated increase of the memory / naïve T-cell ratio is attenuated in C/EBPβ^ΔuORF^ mice. The ratio between CD44^high^ memory T cells and CD44^low^/CD62L^high^ naïve T cells in blood is shown for young (5 months) and old (female 20 months, male 22 months) (A, B) females and (C, D) males for both (a, c) CD8^+^ cytotoxic and (B, D) CD4^+^ helper T cells as was determined by flow cytometry (wt females, n=10 young, n=12 old; wt males n=12 young and old; C/EBPβ^ΔuORF^ females, n=10 young, n=11 old; C/EBPβ^ΔuORF^ males, n=12 young and old). P-values were determined by Student’s t-test, *p<0.05; ***p<0.001.

Aging is associated with a significant decline in motor coordination and muscle strength (Barreto, Huang, & Giffard, 2010; Demontis, Piccirillo, Goldberg, & Perrimon, 2013). In the rotarod test the time is measured that mice endure on a turning and accelerating rod as an indication for their motor-coordination. As expected, rotarod performance decreased with age both for wt female and male mice (**Figure 6A**). Remarkably, rotarod performance was completely preserved in old C/EBPβ^ΔuORF^ females but not in C/EBPβ^ΔuORF^ males. In the beam walking test the required crossing time and number of paw slips of mice traversing a narrow beam are measured. Old mice needed more time to cross the beam reflecting loss of motor coordination upon ageing (**Figure 6B**). The aging-associated increase of the crossing time was less severe in C/EBPβ^ΔuORF^ males and females, although statistically significant only in males (**Figure 6B**). Nevertheless, the strong increase in the number of paw slips in old wt mice is almost completely attenuated in C/EBPβ^ΔuORF^ males and females (**Figure 6C**). Note that the number of paw slips by young C/EBPβ^ΔuORF^ males is already significantly lower compared to young wt males. During the wire hang test the time is measured that mice endure to hang from an elevated wire which serves as an indication for limb skeletal muscle strength (Brooks & Dunnett, 2009). Similar to the rotarod test, the decline in wire hang performance that is seen in old wt mice is completely restored for the female but not for the male C/EBPβ^ΔuORF^ mice (**Figure 6D**).

**Figure 6.**
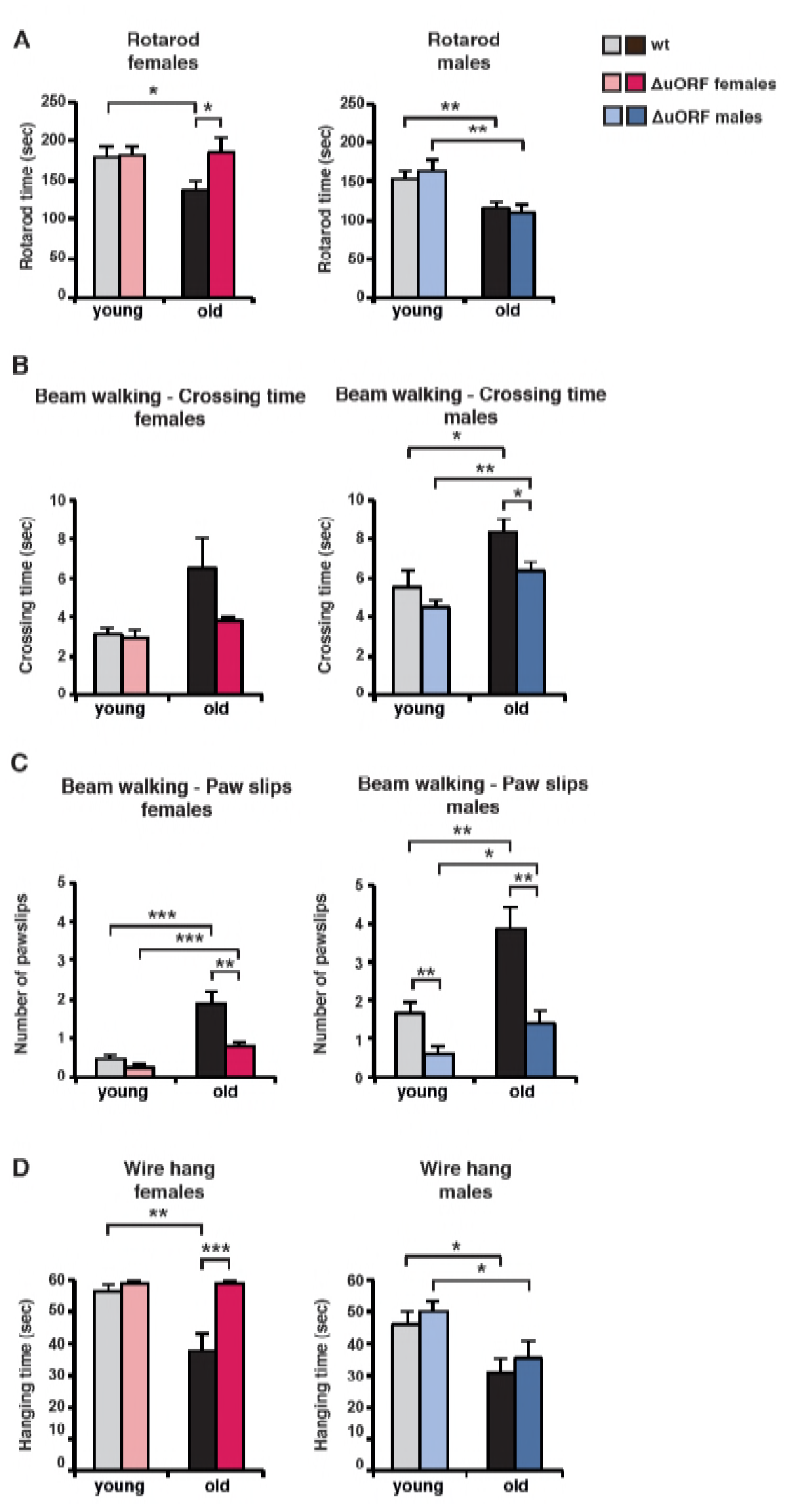
Ageing associated loss of motor coordination and grip strength is attenuated in C/EBPβ^ΔuORF^ mice. (A) Rotarod performance (time in sec of stay on the rotarod) of young (4 months) and old (female 19 months, male 21 months) wt and C/EBPβ^ΔuORF^ mice is shown separately for females (left) and males (right) (wt females, n=11 young, n=12 old; wt males, n=12 young and old; C/EBPβ^ΔuORF^ females, n=11 young, n=12 old; C/EBPβ^ΔuORF^ males, n=12 young, n=11 old). (B) The crossing time (sec) of the beam walking test of young and old wt and C/EBPβ^ΔuORF^ mice, and (C) the number of mistakes (paw slips) made while crossing the beam is shown separately for females and males (wt females, n=11 young, n=12 old; wt males, n=12 young and old; C/EBPβ^ΔuORF^ females, n=11 young, n=12 old; C/EBPβ^ΔuORF^ males, n=12 young, n=10 old). (D) Grip strength as determined with the wire hang test as hanging time (sec) of young and old wt and C/EBPβ^ΔuORF^ mice for females and males separately. N=11 for young wt and C/EBPβ^ΔuORF^ females and for old C/EBPβ^ΔuORF^ females and n= 12 for old wt females; n=11 for young and old wt males; n=12 for young C/EBPβ^ΔuORF^ males and n=10 for old C/EBPβ^ΔuORF^ males. P-values were determined by Student’s t-test, *p<0.05; **p<0.01; ***p<0.001.

Taken together these data demonstrate that the decline in motor coordination and muscle strength is less severe and partly abrogated in female C/EBPβ^ΔuORF^ mice. The results for the old male C/EBPβ^ΔuORF^ mice are not that clear since they show an improved performance only in the beam walking test. One possible explanation is that only the beam walking test measures purely motor coordination skills whereas the results from the rotarod and wire hang tests are influenced in addition by muscle strength and endurance. Old C/EBPβ^ΔuORF^ males thus might have maintained their motor coordination upon ageing but still suffer from an ageing-dependent loss of muscle strength.

By histological examination of different tissues we observed a reduction in some age-related alterations in C/EBPβ^ΔuORF^ mice compared to old wt controls (**Table supplement 3**). We observed a reduced severity of hepatocellular vacuolation and cytoplasmic nuclear inclusions in male C/EBPβ^ΔuORF^ mice; in the pancreas both male and female C/EBPβ^ΔuORF^ mice showed a reduced occurrence and severity of islet cell hyperplasia; in skeletal muscle the number of regenerating muscle fibres was higher in male C/EBPβ^ΔuORF^ mice; the incidence of dermal inflammation was lower in female C/EBPβ^ΔuORF^ mice. Unexpectedly, a slightly increased level of inflammation was detected in the livers of female C/EBPβ^ΔuORF^ mice; a condition believed to be detrimental for glucose homeostasis. However, recently this view was challenged by showing that hepatic inflammation, involving the activation of IKKβ, is beneficial for maintaining glucose homeostasis (Liu et al., 2016). The incidence of other potential age-related pathologies like focal acinar cell atrophy and inflammation in the pancreas, liver polyploidy, spleen lymphoid hyperplasia and extramedulary haematopoiesis, intramuscular adipose tissue infiltration, subcutaneous fat atrophy and bone density were not significantly altered between old wt and C/EBPβ^ΔuORF^ mice. We found slightly reduced plasma IGF-1 levels in old C/EBPβ^ΔuORF^ females compared to old wt females (**Table supplement 3**). A reduction in circulating IGF-1 levels was also found in mice under CR and is believed to be an important mediator of health‐ and lifespan extending effects of CR (Breese, Ingram, & Sonntag, 1991; Mitchell et al., 2016). Taken together our data show that multiple, but not all, ageing associated alterations are attenuated in C/EBPβ^ΔuORF^ mice, particularly in females.

Finally, we performed a comparative transcriptome analysis from livers of 5 and 20 months old wt and C/EBPβ^ΔuORF^ female mice (Müller, 2018). A principal component analysis revealed that there was a clear effect of the genotype on gene expression only in the old mice suggesting that the differences in gene expression between wt and C/EBPβ^ΔuORF^ mice are aging dependent (**figure supplement 6**). This is supported by the finding that in young mice only 42 genes were differentially regulated between wt and C/EBPβ^ΔuORF^ mice (FDR < 0.01; 24 genes upregulated and 18 genes down-regulated in C/EBPβ^ΔuORF^ mice compared to wt mice) while in old mice we found 152 differentially regulated genes (FDR < 0.01; 127 genes upregulated and 25 genes downregulated in C/EBPβ^ΔuORF^ mice compared to wt mice). Gene ontology (GO) analysis using the David database (Huang da, Sherman, & Lempicki, 2009) of the genes upregulated in old C/EBPβ^ΔuORF^ mice in comparison to old wt mice revealed GO terms including “External side of plasma membrane”, “Positive regulation of T-cell proliferation”, and “immune response” (see **Table supplement 4** for the complete list of GO-terms) whereas the GO-terms: “Acute phase” and “Extracellular space” were significantly downregulated (**Table supplement 5**). Acute phase response genes are associated with inflammation and their expression in the liver increases upon ageing (Lee et al., 2012). Like in old C/EBPβ^ΔuORF^ mice expression of acute phase response genes is also inhibited by caloric restriction or treatment with the CR mimetic metformin (Martin-Montalvo et al., 2013) suggesting similar protective mechanisms. The fact that we see the up-regulation of several GO-terms connected to lymphocyte biology fits to the increase in lymphoplasmatic inflammation in the liver of old female C/EBPβ^ΔuORF^ mice observed by pathological analysis (**Table supplement 3**). Although it is generally believed that ageing associated lymphocyte infiltration rather promotes the ageing process by increasing inflammatory signals (Singh et al., 2008), it could also contribute to the removal of senescent or pro-tumorigenic cells, thereby acting protective (Kang et al., 2011). Despite the improved metabolic phenotype of C/EBPβ^ΔuORF^ mice (Zidek et al., 2015), the analysis did not reveal GO-terms related to metabolism. We reasoned that metabolic genes might not be detected as differentially regulated because they are subject of expression heterogeneity in old mice. Comparison between the coefficient of variation of individual transcripts between young and old mice revealed that inter-individual variation of gene expression increases with age in both genotypes (**Figure 7A, B**) supporting earlier observations made by others (White et al., 2015). Direct comparison between old wt and C/EBPβ^ΔuORF^ mice showed that this effect is less pronounced in C/EBPβ^ΔuORF^ mice (**Figure 7C**). KEGG (Kyoto Encyclopedia of Genes and Genomes) pathway and GO-term enrichment analysis of the highly variably expressed genes in the aged livers revealed that in wt mice particularly metabolic genes related to fatty acid metabolism and oxidative phosphorylation were affected which was not observed in C/EBPβ^ΔuORF^ mice (**Figure 7D - Table supplement 6 and 7**). On the other hand genes involved in cell cycle, transcription and RNA biology showed higher inter-individual variation in old C/EBPβ^ΔuORF^ mice compared to wt controls (**Table supplement 7**). These findings suggest that expression control of metabolic genes stays more robust upon aging in C/EBPβ^ΔuORF^ mice. Whether the increased inter-individual variation of metabolic transcripts in old wt mice is a direct effect of the observed increase of the inhibitory-acting LIP isoform or is due to unknown secondary effects has to be clarified in future studies. It is however conceivable that increased transcriptional robustness in the old C/EBPβ^ΔuORF^ mice contributes to the extension in health‐ and lifespan of the female C/EBPβ^ΔuORF^ mice. This idea is supported by our finding that also genes involved in ageing-associated diseases like Nonalcoholic fatty liver disease, Alzheimer’s disease, Parkinson’s disease, Huntington’s disease and cancer (Chemical carcinogenesis) are affected by high inter-individual variation in expression levels in wt but not in C/EBPβ^ΔuORF^ mice (**Figure 7D**).

**Figure 7.**
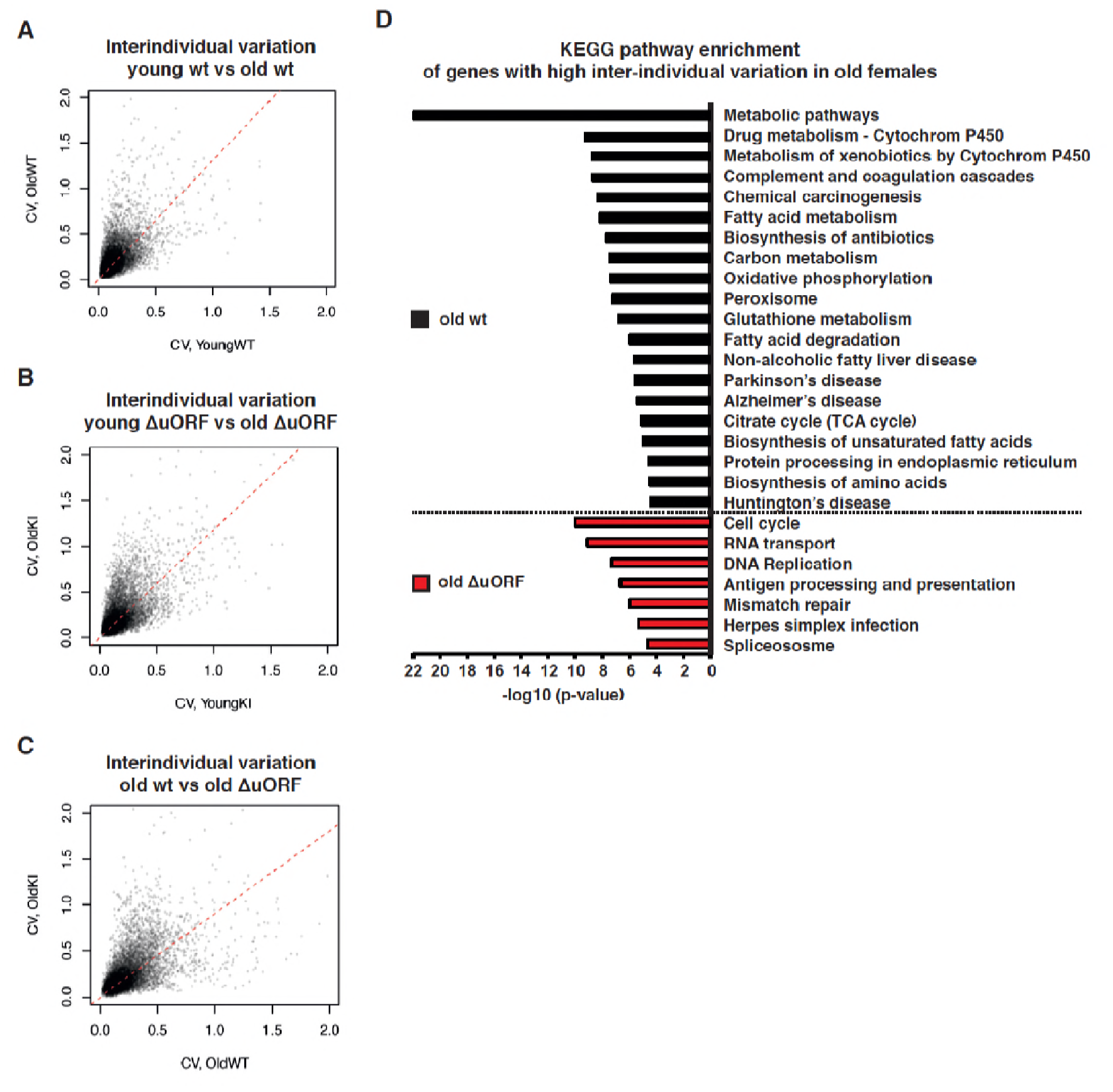
Ageing-associated increase of inter-individual variation of gene expression affects different genes in livers from wt and C/EBPβ^ΔuORF^ mice. (A-C) Inter-individual variability of liver transcripts compared between (A) young (5 months) versus old (20 months) female wt mice, (B) young (5 months) versus old (20 months) C/EBPβ^ΔuORF^ female mice and (C) old wt (20 months) versus old C/EBPβ^ΔuORF^ (20 months) female mice (n=6 for young and old wt and C/EBPβ^ΔuORF^ for A,B,C). Coefficient of variation of transcripts with mean expression >1 FPM is plotted against the coefficient of variation of the other group as indicated. Dashed red line represents linear regression and is shifted towards the side that shows higher inter-individual variability. (D) KEGG pathway enrichment analysis of genes that show increased inter-individual variability in livers from old wt females compared to old C/EBPβΔuORF females (Coefficient of variation of wt genes is more than twice as the coefficient of variation of the same gene in C/EBPβ^ΔuORF^ females) as indicated by the black bars or of genes that show increased inter-individual variability in livers from old C/EBPβ^ΔuORF^ females compared to old wt C/EBPβ^ΔuORF^ females (Coefficient of variation of C/EBPβ^ΔuORF^ genes is more than twice as the coefficient of variation of the same gene in wt females) as indicated by the red bars. The x-axis indicates the p-value. Only pathways that show significant enrichment (FDR < 0.05) are shown.

In summary, reduced signalling through the mTORC1 pathway is thought to mediate many of the beneficial effects of CR or rapamycin treatment (Johnson et al., 2013), and both conditions restrict mTORC1-controlled translation into LIP (Calkhoven et al., 2000; Zidek et al., 2015). These and other studies firmly place LIP function downstream of mTORC1 at the nexus of nutrient signalling and metabolic gene regulation. However, upon ageing, LIP expression increases (the LAP/LIP ratio decreases) in the liver whereas no significant changes in mTORC1 were detected (**figure supplement 7**). Possibly, other pathways play a role in age-related upregulation of LIP as has been described for CUGBP1 (Karagiannides et al., 2001; Timchenko et al., 2006).

Experimental reduction of the transcription factor C/EBPβ-LIP in mice recapitulates many of the effects of CR or treatment with rapamycin, including the reduced cancer incidence and the generally more pronounced extension of lifespan in females (**Figure 8**). We have developed a high-throughput screening screening strategy that allows for discovery of small molecular compounds that suppress the translation into LIP (Zaini et al., 2017). The identification of such compounds or conditions that reduce LIP translation may reveal new ways of CR-mimetic based therapeutic strategies beyond those using mTORC1 inhibition.

**Figure 8.**
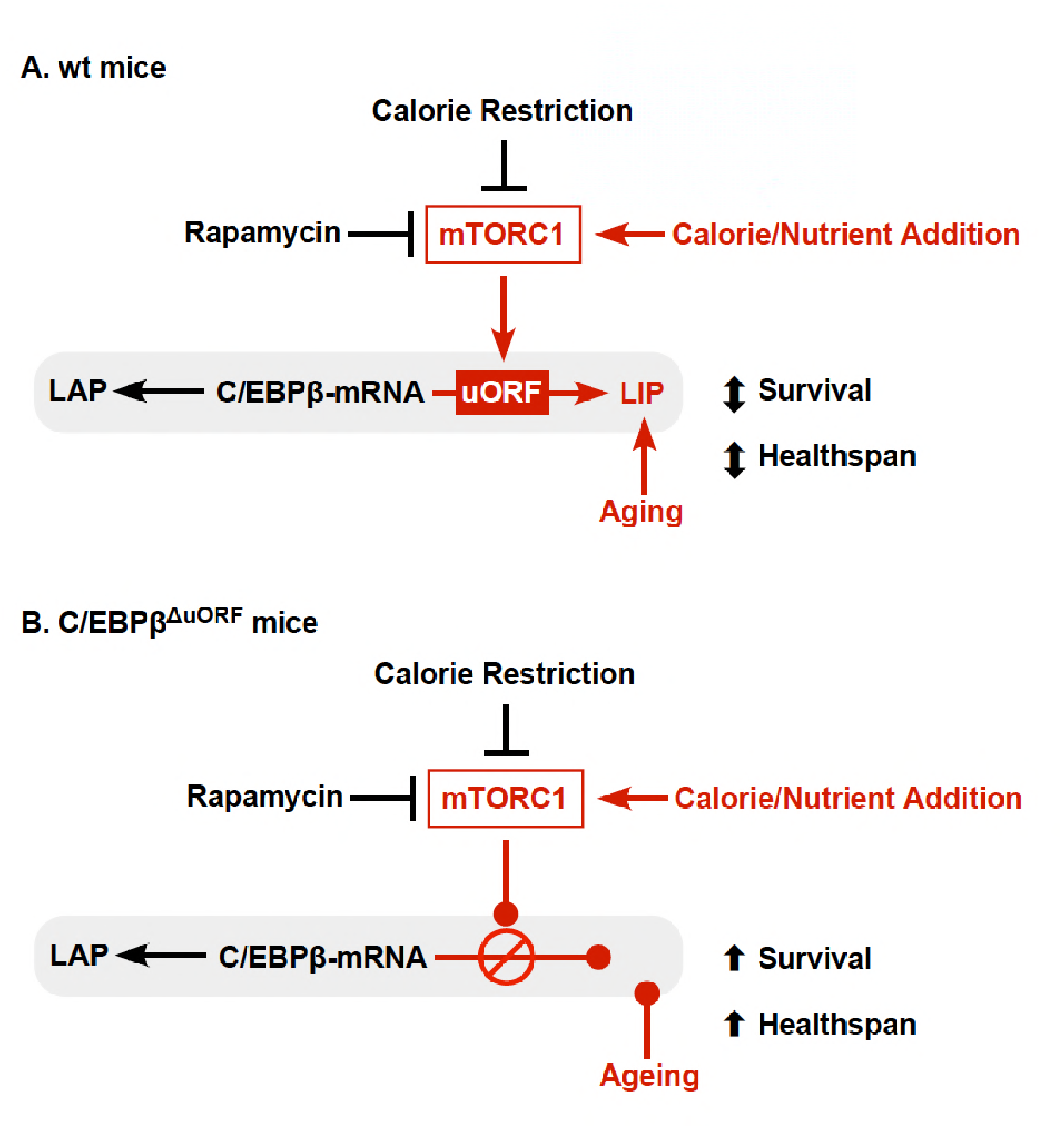
Model explaining regulation of LIP under control of mTORC1. (A) In wt mice C/EBPβ-mRNA translation into LIP is modulated by calorie/nutrient availability through mTORC1 signalling. In addition LIP is upregulated by mechanisms during aging that are not well understood. Expression of LAP is not affected. (B) Deficient expression of LIP in C/EBPβ^ΔuORF^ mice mimics reduced mTORC1 signalling at the level of C/EBPβ function and prevents the age-associated upregulation of LIP and results in health‐ and lifespan extension.

## Material and methods

### Mice

C/EBPβ^ΔuORF^ mice described in (Wethmar et al., 2010) were back-crossed for 12 generations into the C57BL/6J genetic background. Mice were kept at a standard 12-h light/dark cycle at 22°C in individually ventilated cages (IVC) in a specific-pathogen-free (SPF) animal facility on a standard mouse diet (Harlan Teklad 2916). Mice of the ageing cohort were analysed between 3 and 5 months of age (young) and between 18 and 20 months (old females) or between 20 and 22 months (old males) and were derived from the same breeding pairs as mice used in the lifespan experiment. The body weight of the ageing cohorts was determined before the start of the experimental analysis. All of the animals were handled according to approved institutional animal care and use committee (IACUC) protocols of the Thüringer Landesamt für Verbraucherschutz (#03-005/13) and University of Groningen (#6996A).

### Lifespan experiment

C/EBPβ^ΔuORF^ and wt littermates (50 mice from each genotype and gender) derived from mating between heterozygous males and females were subjected to a lifespan experiment. Mice were housed in groups with maximum five female mice or four male mice per cage (separated in genotypes and genders) and did not participate in other experiments. Mice were checked daily and the lifespan of every mouse (days) was recorded. Mice were euthanized when the condition of the animal was judged as moribund and/or to be incompatible with continued survival due to severe discomfort based on the independent assessment of experienced animal caretakers. All mice that were found dead or were euthanized underwent necropsy with a few exceptions when the grade of decomposition of dead animals prevented further examination (number of mice without necropsy: n=0 for wt females; n=2 for C/EBPβ^ΔuORF^ females; n=3 for wt males and n=5 for C/EBPβ^ΔuORF^ males. Survival curves were calculated with the Kaplan-Meier method. Statistical significance was determined by the log-rank test using GraphPath Prism 6 software. Maximum lifespan was determined by the number of mice for each genotype that were within the 10% longest-lived mice of the combined (wt and C C/EBPβ^ΔuORF^) cohorts. Statistical significance of observed differences was calculated with Fisher’s exact test. In addition, the mean lifespan (±SEM) of the 10% longest lived mice within one genotype was compared to the mean lifespan of the 10% longest lived mice of the other genotype, and the statistical significance was calculated with the Student’s T-test.

### Tumour incidence

Suspected tumour tissue found during necropsy of the lifespan cohorts was fixed in 4% paraformaldehyde and Haematoxylin & Eosin stained tissue slices were analysed by experienced board-certified veterinary pathologists of the Dutch Molecular Pathology Centre (Utrecht University) to diagnose the tumour type. Tumour incidence was calculated as percentage of mice with pathologically confirmed tumours in respect to all mice from the same cohort that underwent necropsy. Tumour occurrence was defined as the time of death of an animal in which a pathologically confirmed tumour was found. Tumour load was defined as number of different tumour types found in the same mouse and tumour spread was defined as number of different organs harbouring a tumour within the same mouse irrespective of the tumour type with the exception that in those cases in which different tumour types were found in the same organ a number >1 was rated.

### Motor coordination experiments

Rotarod test: Mice were habituated to the test situation by placing them on a rotarod (Ugo Basile) with constant rotation (5 rpm) for 5 min at two consecutive days with two trials per mouse per day separated by an interval of 30 min. In the test phase two trials per mouse were performed with accelerating rotation (2-50 rpm within 4 min) with a maximum trial duration of 5 min in which the time was measured until mice fell off the rod. Beam walking test: Mice were trained by using a beam of 3 cm width and 100 cm in length at two consecutive days (one trial per mouse per day). At the test day mice had to pass a 1 cm wide beam, 100 cm in length and beam crossing time and number off paw slips upon crossing was measured during 3 trials per mouse that were separated by an interval of 20 min. To determine of the number of mistakes the number of paw slips per trial was counted upon examination of recorded videos. Wire Hang test: To measure limb grip strength mice were placed with their four limbs at a grid with wire diameter of 1 mm at 20 cm over the layer of bedding material and the hanging time was measured until mice loosened their grip and fell down. Three trials of maximal 60s per mouse were performed that were separated by an interval of 30 min.

### Body composition

The body composition was measured using an Aloka LaTheta Laboratory Computed Tomograph LCT-100A (Zinsser Analytic) as described in (Zidek et al., 2015). Percentage body fat was calculated in relation to the sum of lean mass and fat mass.

### Bone measurements

Bones of the hind legs were freed from soft tissue and fixed in 4% paraformaldehyde. For determination of the bone volume, trabecular thickness, trabecular number and trabecular separation femurs were analysed by micro CT (Skyscan 1176, Bruker) equipped with an X-ray tube (50 kV/500 mA). The resolution was 9 mm, rotation step was set at 1°C, and a 0.5 mm aluminium filter was used. For reconstruction of femora, the region of interest was defined 0.45 mm (for trabecular bone) or 4.05 mm (for cortical bone) apart from the distal growth plate into the diaphysis spanning 2.7 mm (for trabecular bone) or 1.8 mm (for cortical bone). Trabecular bone volume/tissue volume (%), trabecular number per μm, trabecular thickness (mm) and trabecular separation (intertrabecular distance, mm) was determined according to guidelines by ASBMR Histomorphometry Nomenclature Committee (Dempster et al., 2013).

### Glucose tolerance

The intraperitoneal (i.p.) glucose tolerance test (IPGTT) was performed as described in (Zidek et al., 2015). Mice without initial increase in blood glucose concentration were excluded from the analysis.

### Flow cytometry

Blood cells from 300 μl blood were incubated in RBC-Lysis buffer (Biolegend) to lyse the red blood cells. Remaining cells were washed and incubated with a cocktail of fluorochrome-conjugated antibodies (CD4-PE-Cy7 (#552775) and CD62L-FITC (#561917) from BD Pharmingen; CD3e-PE (#12-0031), CD8a-eFluor 450 (#48-0081) and CD44-APC (#17-0441) from eBioscience.), incubated with propidium iodide for the detection of dead cells and analysed using the FACSCanto II analyser (BD Biosciences). The following T cell subsets were quantified: CD3^+^, CD8^+^, CD44^high^ cytotoxic memory T cells; CD3^+^, CD8^+^, CD44^low^, CD62L^high^ cytotoxic naïve T cells, CD3^+^, CD4^+^, CD44^high^ helper memory T cells and CD3^+^, CD4^+^, CD44^low^, CD62L^high^ helper naïve T cells.

### Histology

Tissue pieces were fixed with 4% paraformaldehyde and embedded in paraffin. Sections were stained with Haematoxylin and Eosin (H&E) and age-related pathologies or tumour types were analysed by experienced board-certified veterinary pathologists of the Dutch Molecular Pathology Centre (Utrecht University). Semi-quantification of muscle regeneration was done by counting the number of myofibers with a row of internalized nuclei (>4) for five 200x fields. Other ageing-associated lesions were scored subjectively and the severity of the lesions was graded on a scale between 0 and 3 with 0 = absent; 1 = mild; 2 = moderate and 3 = severe.

### Immunoblotting and quantification

Mouse liver tissue was homogenized on ice with a glass douncer in RIPA buffer (150 mM NaCl, 1% NP40, 0.5% sodium deoxycholate, 0.1% SDS, 50 mM TRIS pH 8.0 supplemented with protease and phosphatase inhibitors) and sonicated. Equal amounts of total protein were separated by SDS-PAGE, transferred to a PVDF membrane and incubated with the following antibodies: C/EBPβ (E299) and β-actin (ab16039) from Abcam; 4E-BP1 (C-19) from Santa Cruz; phospho-p70S6K (Thr389) (108D2), p70S6K (#9202) and phospho-4E-BP1 (Thr 37/46) (#9459) from Cell Signaling Technology and HRP-linked anti rabbit IgG from GE Healthcare. Lightning Plus ECL reagent (Perkin Elmer) was used for detection and for re-probing membranes were incubated in Restore Western Blot Stripping buffer (Perbio). The detection and quantification of protein bands was performed with the Image Quant LAS 4000 Mini Imager (GE Healthcare) using the supplied software.

### Quantitative real-time PCR

Mouse liver or visceral fat tissue was homogenized on ice with a motor driven pellet pestle (Kontes) in the presence of QIAzol reagent (QIAGEN) and total RNA was isolated as described in(Zidek et al., 2015). cDNA synthesis was performed from 1 μg of total RNA with the Transcriptor First Strand cDNA Synthesis Kit (Roche) using random hexamer primers. qRT-was performed with the LightCycler 480 SYBR Green I Master mix (Roche) using the following primers: β-actin: 5’-AGA GGG AAA TCG TGC GTG AC-3’ and 5’-CAA TAG TGA TGA CCT GGC CGT-3’; C/EBPβ: 5’-CTG CGG GGT TGT TGA TGT-3’ and 5’-ATG CTC GAA ACG GAA AAG GT-3’; CD68: 5’-GCC CAC CAC CAC CAG TCA CG-3’ and 5’- GTG GTC CAG GGT GAG GGC CA-3’.

### Enzyme-linked immune-sorbent assay (ELISA)

Plasma was prepared as described in (Zidek et al., 2015) and the IGF-1 specific ELISA was performed according to the instructions of the manufacturer (BioCat).

### RNA-seq Analysis

Liver tissue from young (5 months) and old (20 months) wt and C/EBPβ^ΔuORF^ mice (from six individuals per group) was homogenized on ice with a motor driven pellet pestle (Kontes) in the presence of QIAzol reagent (Qiagen) and total RNA was isolated as described in Zidek et al(Zidek et al., 2015). Preparation of the sequencing libraries was performed using the TruSeq Sample Preparation V2 Kit (Illumina) according to the manufacturer’s instructions. High-throughput single-end sequencing (65 bp) of the libraries was performed with an Illumina HiSeq 2500 instrument. Reads were aligned and quantified using STAR 2.5.2b(Dobin et al., 2013) against primary assembly GRCm38 using Ensembl gene build 86 (http://www.ensembl.org). Genes with average expression level below 1 fragment per million (FPM) were excluded from the analysis. A generalized linear model was used to identify differential gene expression using EdgeR package (McCarthy, Roche, & Forde, 2012; Robinson, McCarthy, & Smyth, 2010). The library normalization was left at the standard setting (trimmed mean of M-values, TMM). The resulting p-values were corrected for multiple testing using the Benjamini-Hochberg procedure. Data visualization, calculation of CV (coefficient of variation) and statistical tests were conducted using custom R scripts (de Jong, 2018). Gene ontology (GO) analysis was performed using the DAVID database version 6.8 (Huang da et al., 2009) with default DAVID database setting with medium stringency and *Mus musculus* background. KEGG pathway analysis was performed using gProfiler tool (Reimand et al., 2016). For dataset see (Müller, 2018).

### Statistical analysis

Biological replication is indicated (n=x). All graphs show average ± standard error of the mean (s.e.m.). The unpaired, two-tailed Student’s t-Test was used to calculate statistical significance of results with * p< 0.05; ** p<0.01; *** p< 0.001. Significance of the differences in survival curves were analysed using the log-rank test and significance of the difference in maximum lifespan (number of mice from one cohort within the 10% longest lived mice calculated from the combined cohort) and tumour incidence was calculated using the Fisher’s exact test with *p<0.05.

## Acknowledgements

We thank Rafael de Cabo, NIA Baltimore for advice with the lifespan experiment. At the ERIBA/UMCG Groningen we thank Mirjam Koster for technical assistance with histology and Gerald de Haan for critical reading of the manuscript. At the FLI we thank we thank Sabrina Eichwald for technical assistance, Lucien Frappart and Dominique Galendo for advice on necropsy, the staff of the animal house facility in particular Anja Baar and Juliane Brüchert for massive support with the lifespan experiment, Nico Andreas for advice concerning the flow cytometry experiments, Anne Gompf for technical assistance with flow cytometry, Maik Baldauf for paraffin embedding and Christina Valkova for advice with the motor coordination experiments. L.M.Z was supported by the Deutsche Forschungsgemeinschaft (DFG) through a grant to C.F.C. (CA 283/1-1) and a grant to J.V.M. (MA-3975/2-1). T.A. was supported by the Leibniz Graduate School on Ageing and Age-Related Diseases (LGSA; www.flileibniz.de/phd) P.L. was supported by the Collaborative Research Centre 1149 ‘Trauma’ (INST 40/492-1) and DFG priority program Immunobone (Tu220/6) to a grant to J.P.T.

## Author contributions

C.M., M.Z, T.A, P.L, V.K, M.A.Z, G.K, and J.M. performed experiments; L.H, A. de B. performed patho-histological analysis; T.de J, V.G. performed bioinformatic analysis; J.P.T, D.V, J.von M. and Z-Q. W. designed and advised on animal experiments. C.M., L.M.Z., J. von M., V.G., Z-Q. W and C.F.C wrote the manuscript. C.F.C supervised the project.

---

**Figure supplement 1.**
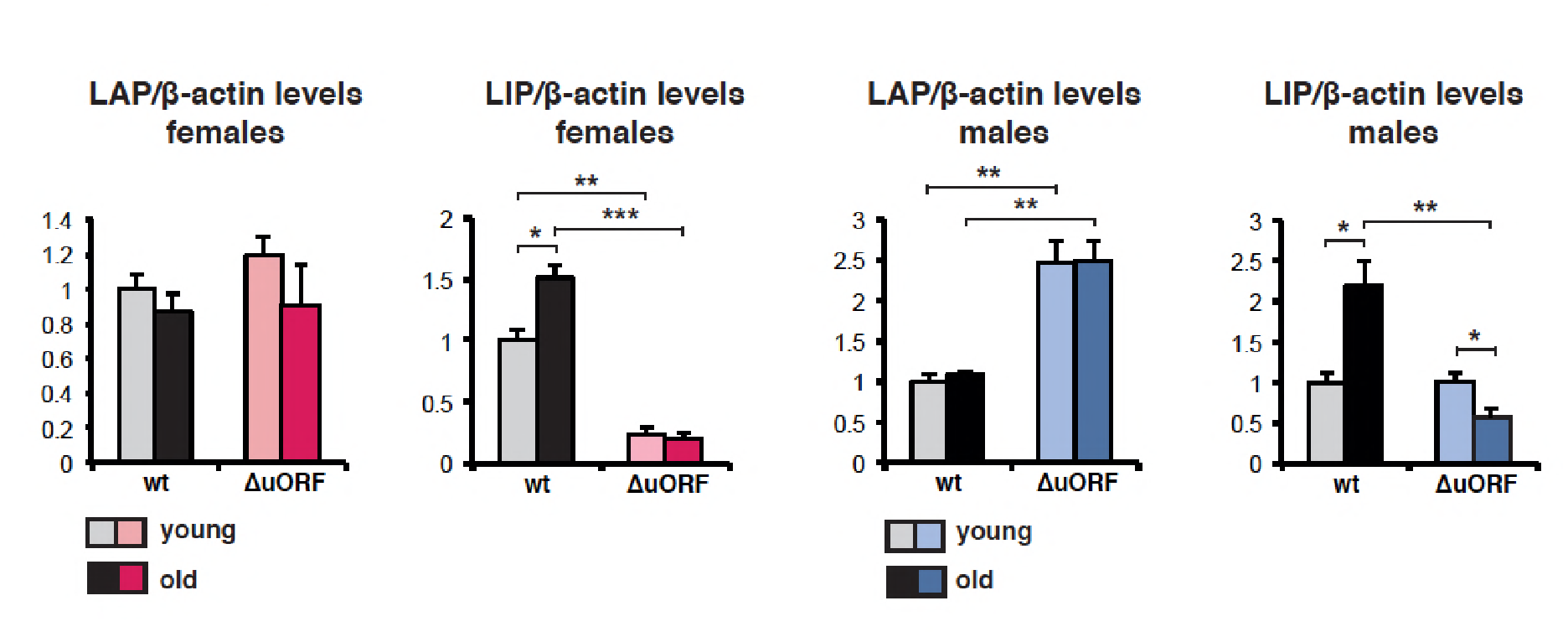
Quantification chemiluminescence digital imaging of LAP or LIP signals in livers in respect to β-actin levels using immunoblots presented in Figure 1A and B and additional immunoblots from young (5 month) and old (female 20 months, male 22 months) mice (wt females, n=9 young and old; wt males, n=10 young, n=11 old; C/EBPβ^ΔuORF^ females, n=10 young, n=11 old; C/EBPβ^ΔuORF^ males, n=11 young, n=10 old). P-values were determined by Student’s t-test, *p<0.05; **p<0.01; ***p<0.001.

**Figure supplement 2.**
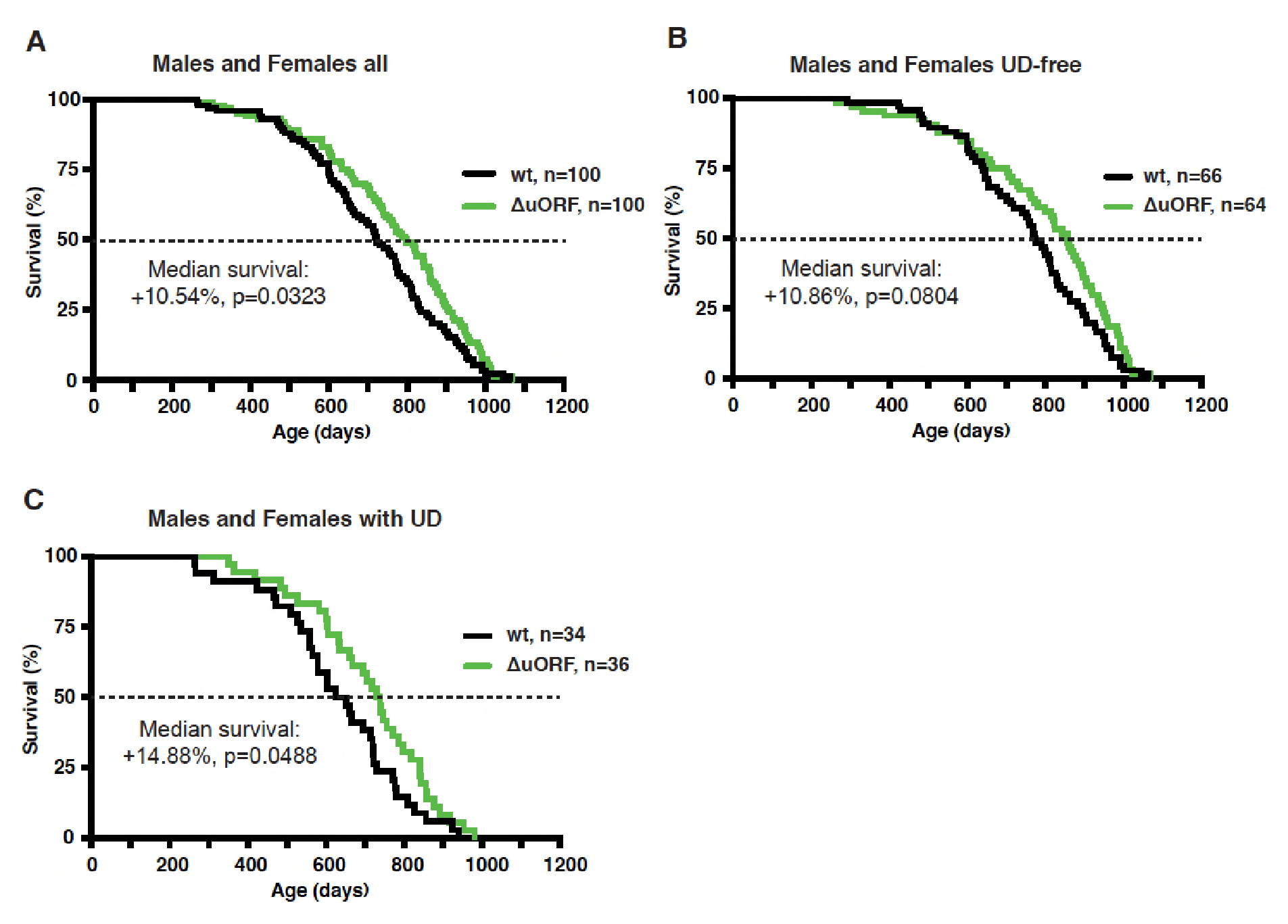
Survival curves of wt and C/EBPβ^ΔuORF^ mice with females and males combined (A) the complete cohorts, (B) UD-free mice cohorts and (C) cohorts of mice with UD. For survival curves the increase in median survival (%) of the C/EBPβ^ΔuORF^ compared to wt mice and statistical significance of the increase in the overall survival as determined by the log-rank test is indicated in the figure.

**Figure supplement 3.**
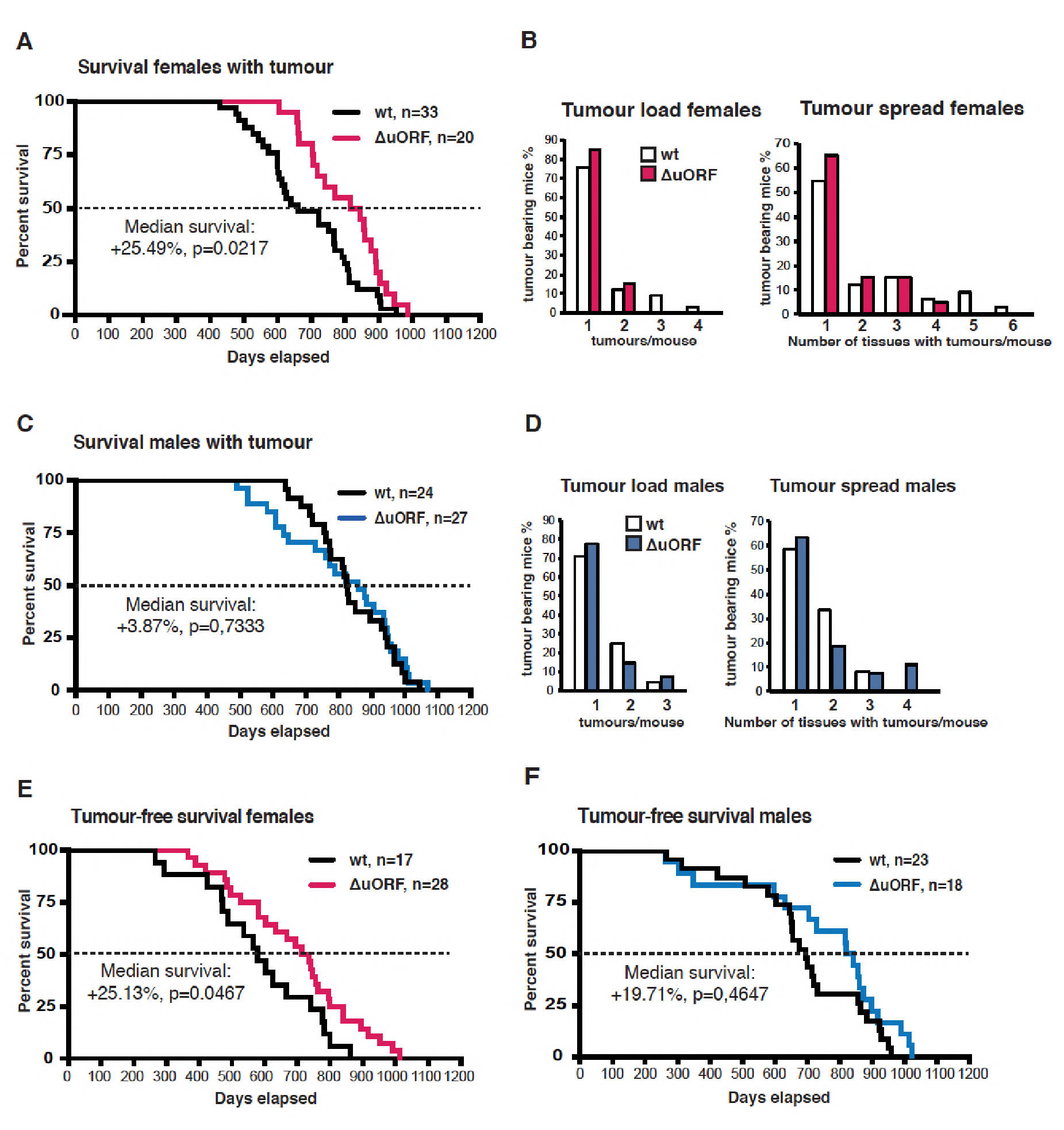
Survival curves of wt and C/EBPβ^ΔuORF^ tumour-bearing female mice. (B) At the left, tumour load in wt and C/EBPβ^ΔuORF^ females as determined by the number of different tumour types found per tumour bearing mouse. At the right, tumour spread in wt and C/EBPβ^ΔuORF^ females as determined by the number of tissues affected by tumours in tumour bearing mice irrespective of the tumour type (in case different tumour types were found in the same tissue it was counted as >1). (C) Survival curves of wt and C/EBPβ^ΔuORF^ tumour-bearing male mice. (D) At the left tumour load and at the right tumour spread wt and C/EBPβ^ΔuORF^ males as described under (C). (E) Survival curves of wt and C/EBPβ^ΔuORF^ tumour-free female mice. (F) Survival curves of wt and C/EBPβ^ΔuORF^ tumour-free male mice. For survival curves the increase in median survival (%) of the C/EBPβ^ΔuORF^ compared to wt mice and statistical significance of the increase in the overall survival as determined by the log-rank test is indicated in the figure.

**Figure supplement 4.**
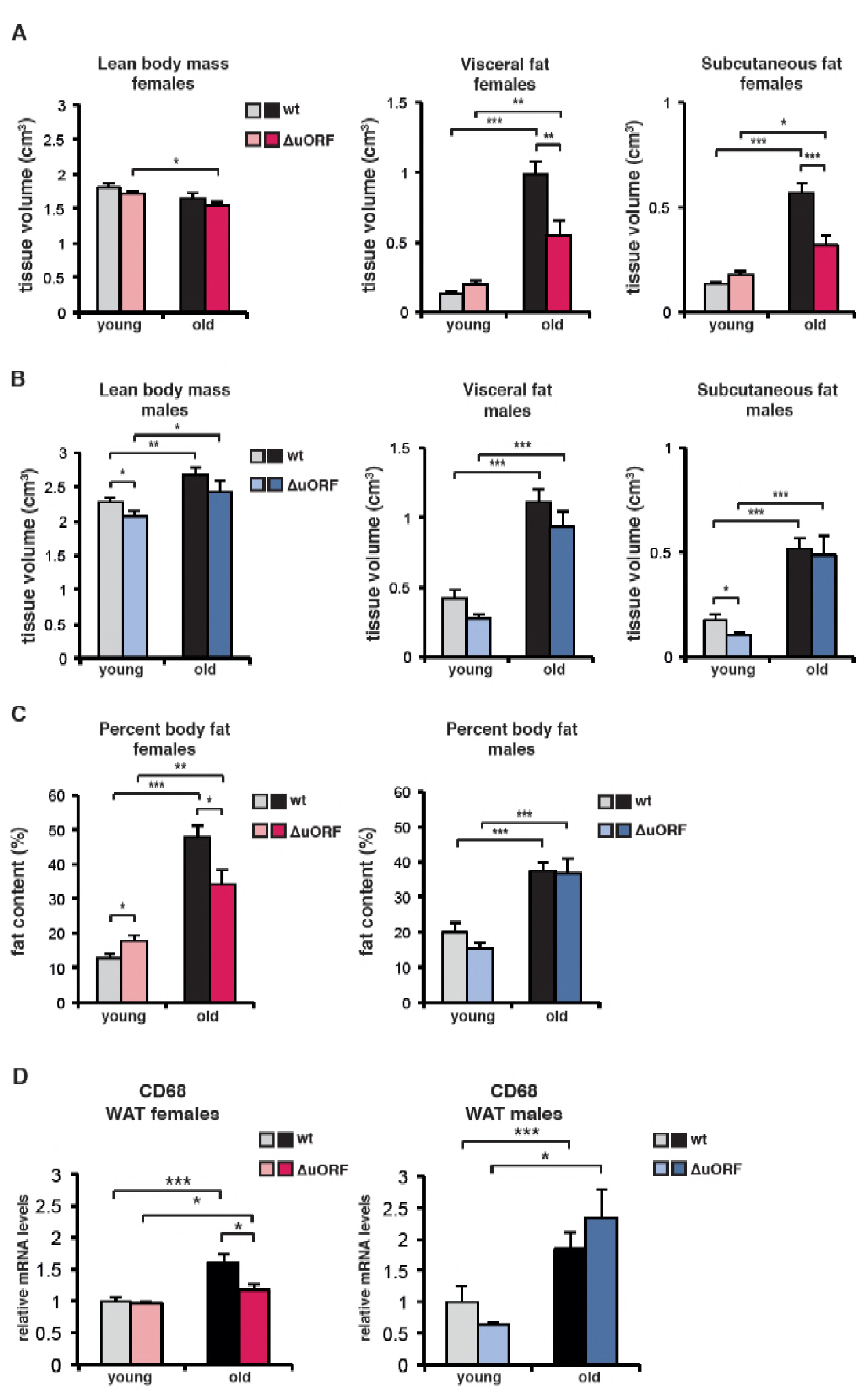
Body composition of young (4 months) and old (females 19 months, males 21 months) wt and C/EBPβ^ΔuORF^ mice was determined by computer tomography (CT) analysis. (A) The bar graphs show tissue volumes (cm^3^) of lean body mass, visceral fat and subcutaneous fat for females, and (B) for males. (C) Percentage of body fat for females (left) or males (right). (wt females, n=11 young and old; wt males, n=12 young and old; C/EBPβ^ΔuORF^ females, n=9 young, n=11 old; C/EBPβ^ΔuORF^ males, n=11 young, n=9 old). (d) Relative CD68 mRNA levels as determined by quantitative real-time PCR in visceral fat of young (5 months) and old (females 20 months, males 22 months) wt and C/EBPβ^ΔuORF^ females (left) and males (right) as a measure for macrophage infiltration (wt females, n=11 young and old; wt males, n=11 young, n=12 old; C/EBPβ^ΔuORF^ females, n=11 young and old; C/EBPβ^ΔuORF^ males, n=11 young, n=10 old). P-values were determined by Student’s t-test, *p<0.05; **p<0.01; ***p<0.001.

**Figure supplement 5.**
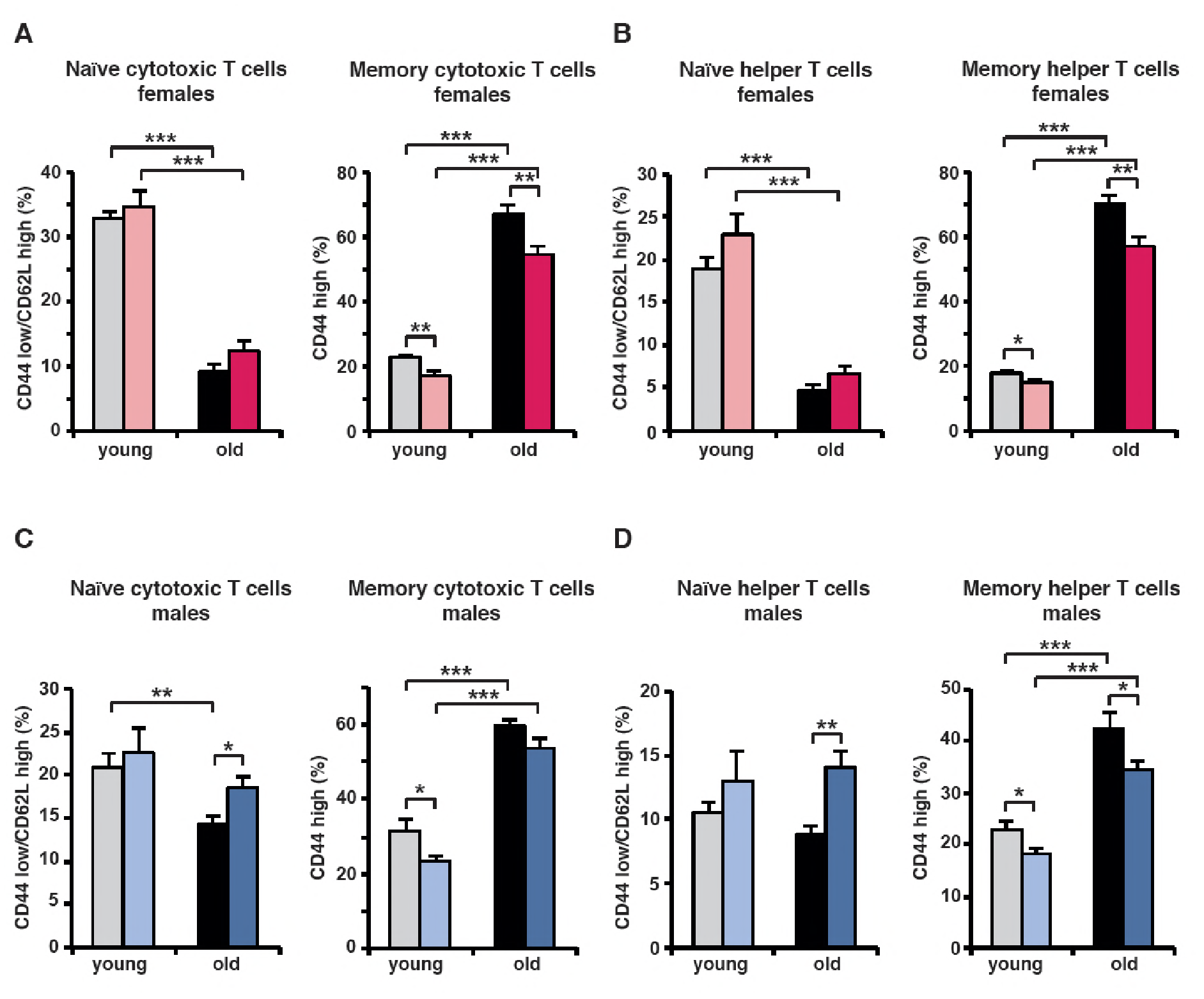
(A) At the left, for females the percentage naïve cytotoxic T cells (CD44^low^/CD62L^high^) in respect to the total amount of cytotoxic T cells (CD8^+^); at the right, percentage of memory (CD44^high^) cytotoxic T cells in respect to the total amount of cytotoxic (CD8^+^) T cells. (B) At the left, for females the percentage naïve helper T cells (CD44^low^/CD62L^high^) in respect to the total amount of helper T cells (CD4^+^); at the right percentage memory T cells (CD44^high^) in respect to the total amount of helper T cells (CD4^+)^. (C) The same analysis as in (A) for males. (D) The same analysis as in (B) for males. (wt females, n=10 young, n=12 old; wt males, n=12 young and old; C/EBPβ^ΔuORF^ females, n=10 young, n=12 old; C/EBPβ^ΔuORF^ males, n=12 young and old). P-values were determined by Student’s t-test, *p<0.05; **p<0.01; ***p<0.001.

**Figure supplement 6.**
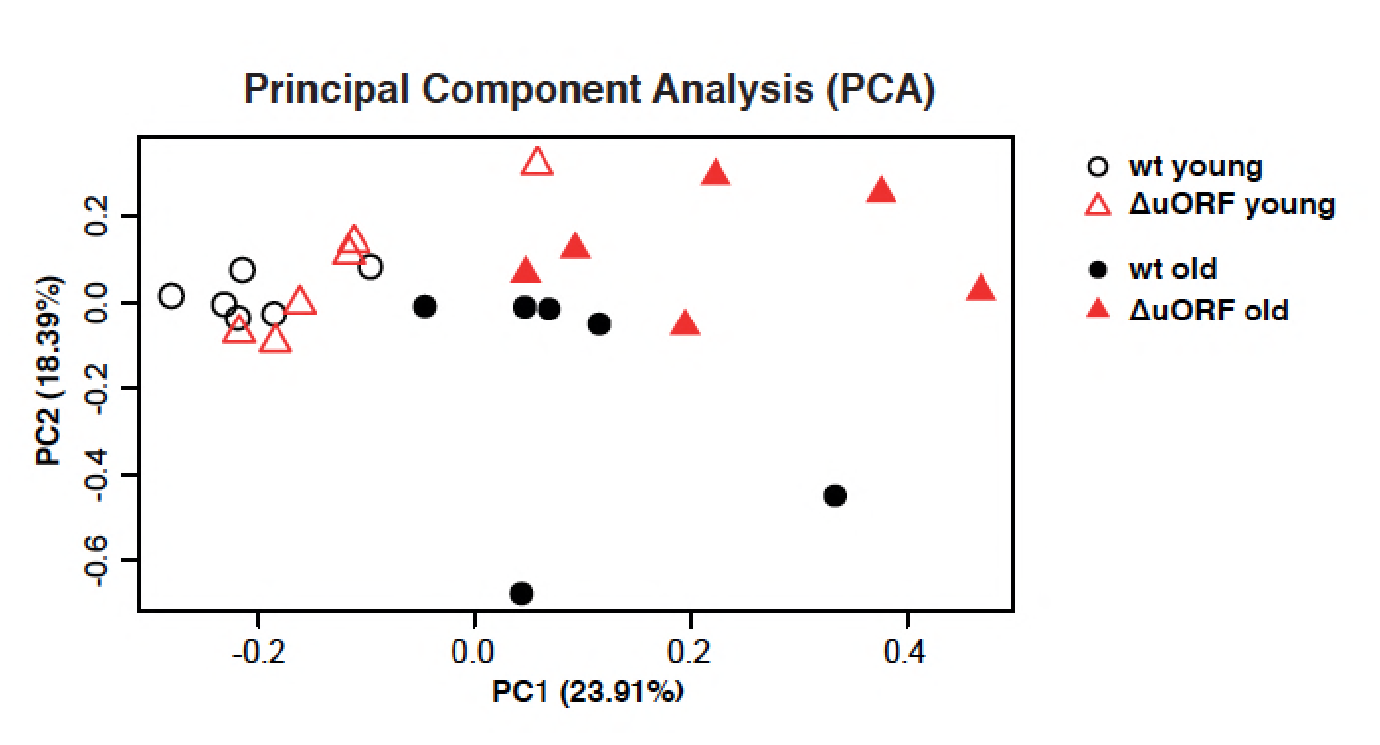
Principal component analysis (PCA) of transcriptome profiles obtained from livers of 5 and 20 months old wt and C/EBPβΔuORF female mice (n=6 for all groups). The main principal component PC1 (x axis, 23,91%) reflects expression changes due to ageing independent of the genotype and the second component PC2 (y axis, 18,39%) reflects expression changes due to the genotype.

**Figure supplement 7.**
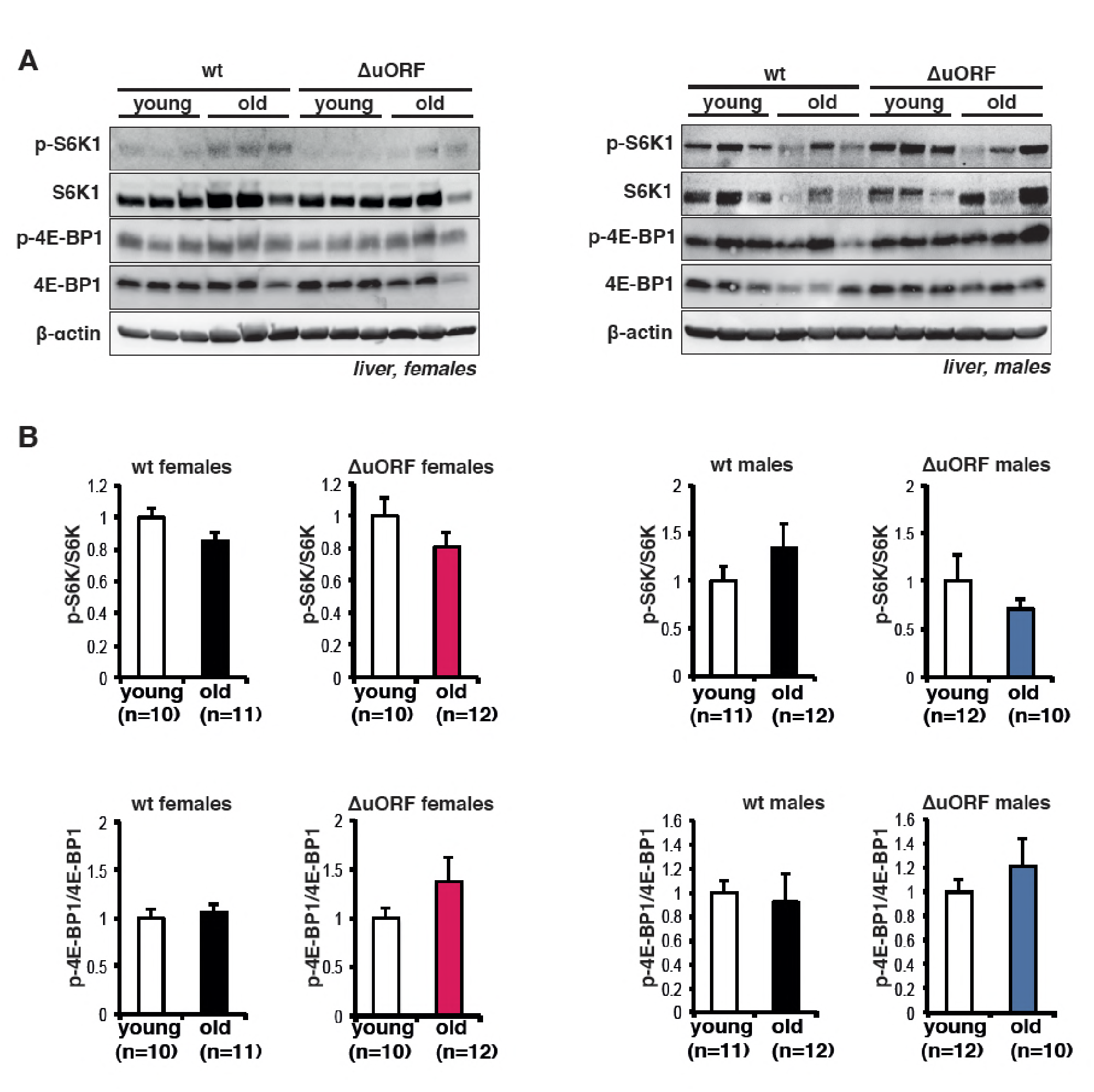
(A) Immunoblots show pan-levels and phosphorylation of the mTORC1 downstream targets S6K1 and 4E-BP1 in liver extracts from young (5 month) and old (females 20 months, males 22 months) of female (left) and male (right) wt and C/EBPβ^ΔuORF^ mice. β-actin detection served as loading control. (B) Quantification by chemiluminescence digital imaging of phosphorylated proteins in respect to the respective pan levels from the blots presented in (A) and from additional immunoblots (wt females, n=10 young, n=11 old; wt males, n=11 young, n=12 old; C/EBPβ^ΔuORF^ females, n=10 young, n=12 old; C/EBPβ^ΔuORF^ males, n=12 young, n=10 old). P-values were determined by Student’s t-test, *p<0.05; **p<0.01; ***p<0.001.

**Table supplement 1.** Lifespan experiment summary of results

| Genotype | sex | N<sup>1</sup> | Median<sup>2</sup> | Increase<sup>3</sup> | P<sub>Surv</sub><sup>4</sup> | Mean<sup>5</sup> | SEM<sup>6</sup> | N<sub>max</sub><sup>7</sup> | P<sub>max</sub><sup>8</sup> |
| --- | --- | --- | --- | --- | --- | --- | --- | --- | --- |
| **All mice** |  |  |  |  |  |  |  |  |  |
| C/EBPβ<sup>ΔuORF</sup> | Females | 50 | 751.5 | 20.63 | 0.0014 | 747.42 | 23.7 | 9 | 0.0157 |
| wt | Females | 50 | 623.0 |  |  | 650.96 | 22.5 | 1 |  |
| C/EBPβ<sup>ΔuORF</sup> | Males | 50 | 833.5 | 5.24 | 0.4647 | 781.90 | 28.8 | 6 | 0.7407 |
| wt | Males | 50 | 792.0 |  |  | 777.68 | 24.8 | 4 |  |
| C/EBPβ<sup>ΔuORF</sup> | Both | 100 | 797.0 | 10.54 | 0.0323 | 764.66 | 18.6 | 13 | 0.2381 |
| wt | Both | 100 | 721.0 |  |  | 714.32 | 17.8 | 7 |  |
| **Mice ulcerative dermatitis (UD)-free** |  |  |  |  |  |  |  |  |  |
| C/EBPβ<sup>ΔuORF</sup> | Females | 24 | 860.5 | 16.28 | 0.0017 | 809.29 | 34.10 | 5 | 0.0755 |
| wt | Females | 31 | 740.0 |  |  | 686.07 | 29.71 | 1 |  |
| C/EBPβ<sup>ΔuORF</sup> | Males | 40 | 857.5 | 3.44 | 0.8282 | 789.10 | 33.11 | 6 | 0.2710 |
| wt | Males | 35 | 829.0 |  |  | 821.49 | 24.75 | 2 |  |
| C/EBPβ<sup>ΔuORF</sup> | Both | 64 | 857.5 | 10.86 | 0.0804 | 796.67 | 24.19 | 8 | 0.3936 |
| wt | Both | 66 | 773.5 |  |  | 757.88 | 20.78 | 5 |  |
| **Mice with UD** |  |  |  |  |  |  |  |  |  |
| C/EBPβ<sup>ΔuORF</sup> | Females | 26 | 711.5 | 22.88 | 0.0122 | 690.31 | 37.37 | 5 | 0.0634 |
| wt | Females | 19 | 579.0 |  |  | 593.68 | 30.65 | 0 |  |
| C/EBPβ<sup>ΔuORF</sup> | Males | 10 | 802.5 | 12.40 | 0.3299 | 753.1 | 58.63 | 1 | 1.0 |
| wt | Males | 15 | 714.0 |  |  | 675.47 | 51.28 | 2 |  |
| C/EBPβ<sup>ΔuORF</sup> | Both | 36 | 733.5 | 14.88 | 0.0488 | 707.75 | 26.67 | 5 | 0.4297 |
| wt | Both | 34 | 638.5 |  |  | 629.77 | 28.78 | 2 |  |

<sup>1</sup>Number of animals in the cohort
<sup>2</sup>Median survival of the cohort (days)
<sup>3</sup>Increase of the median survival (percent)
<sup>4</sup>P-value of the increased survival (log-rank test)
<sup>5</sup>Mean lifespan of the cohort (days)
<sup>6</sup>Standard error of the mean
<sup>7</sup>Number of mice in the cohort in the longest-lived decile of the combined cohort (wt and C/EBPβ<sup>ΔuORF</sup>)
<sup>8</sup>P-value of increased N<sub>max</sub><sup>7</sup> (Fisher's exact test)

**Table supplement 2.** Tumour spectrum in wt and C/EBPβ^ΔuORF^ mice

| Tumour type | Location | wt females | $\Delta uORF$ females | wt males | $\Delta uORF$ males |
| --- | --- | --- | --- | --- | --- |
| Lymphoma * | different | 15 | 6 | 1 | 4 |
| Hepatocellular carcinoma | liver | 8 | 4 | 14 | 6 |
| Histiocytic sarcoma | liver, colon, intestine, spleen | 8 | 3 | 14 | 9 |
| Myeloid neoplasms ** | different | 4 | 4 | 4 | 6 |
| Adenocarcinoma | pituitary, mammary gland, haderian gland, intestine | 2 | 2 | 0 | 3 |
| Bronchio-alveolar carcinoma | lung | 0 | 1 | 0 | 3 |
| Mast cell tumour | stomach, liver | 1 | 0 | 0 | 1 |
| Anaplastic sarcoma | leg, side | 1 | 1 | 0 | 0 |
| Osteosarcoma | leg | 1 | 0 | 0 | 0 |
| Undifferentiated sarcoma | pancreas, colon, kidney | 0 | 0 | 0 | 1 |
| Squamous cell carcinoma | penis | 0 | 0 | 1 | 0 |
| Benign neoplasms | bladder, neck, spleen, liver | 3 | 1 | 1 | 0 |
| Unidentified*** | liver, kidney, colon, spleen, lung, throat, abdominal cavity, adrenal gland | 6 | 5 | 3 | 2 |

\*incl. lymphoid leukaemia
\*\*incl. malignant round cell neoplasms
\*\*\*tumour type could not be unequivocally determined due to inadequate quality of the fixed tumour tissue
Absolute numbers of mice with the indicated tumour type of tumours found during necropsy for wt and C/EBP $\beta^{\Delta uORF}$ males and females are shown. Note that the total number of tumours is higher than the number of tumour-bearing mice due to the eventual occurrence of different tumour types in the same mouse.

**Table supplement 3.**
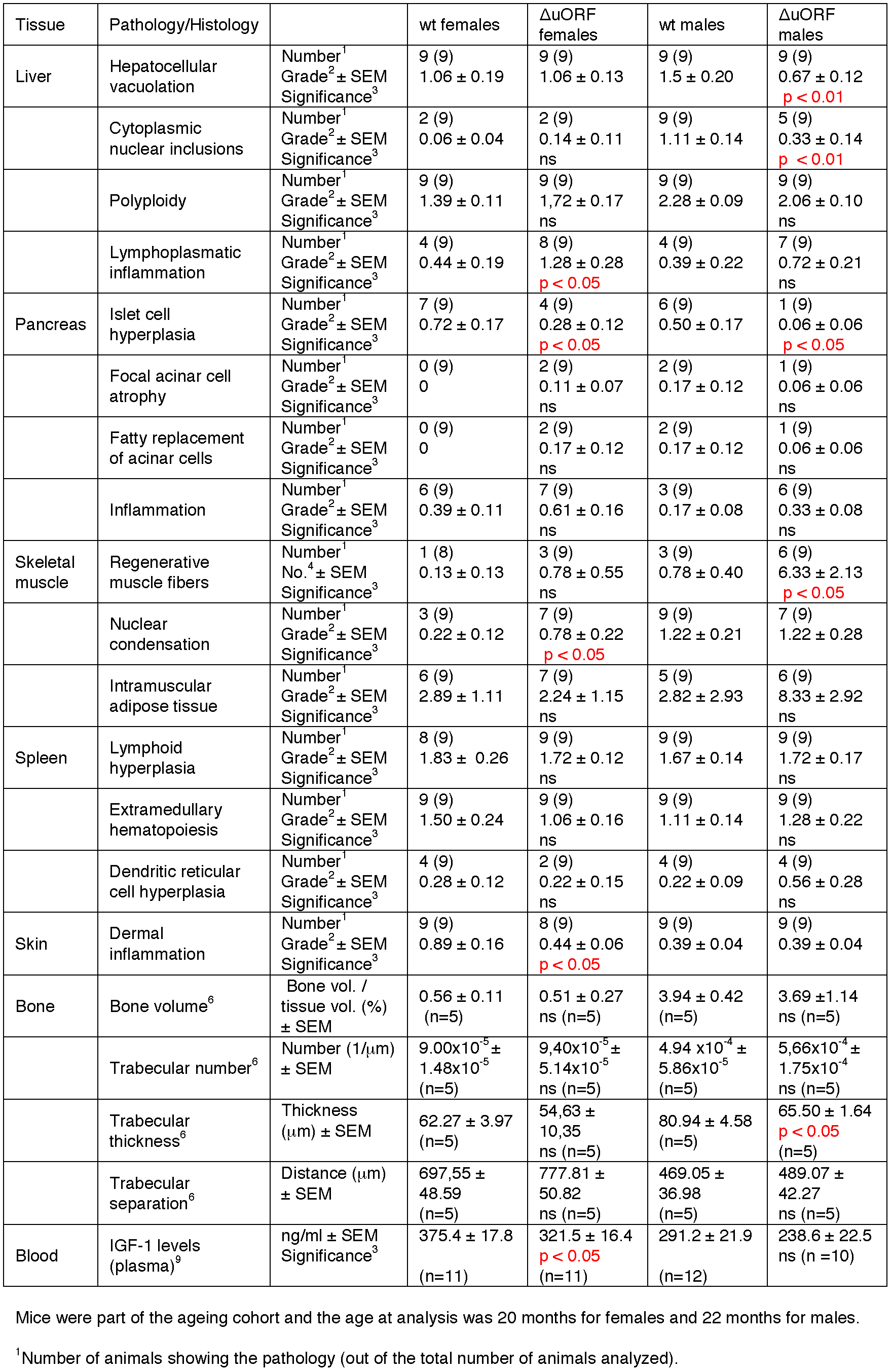
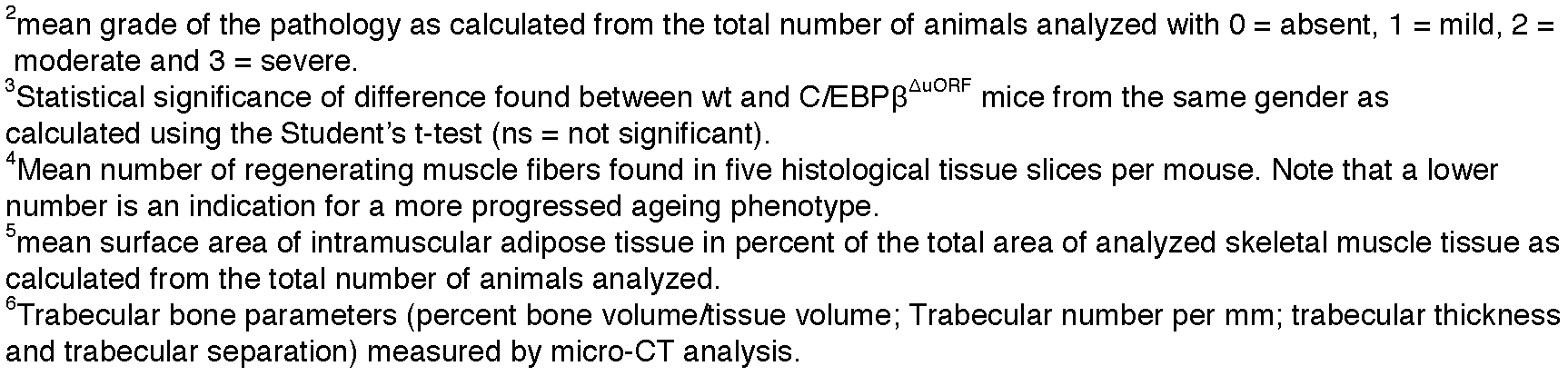
Occurrence of ageing-associated pathologies in wt and C/EBPβ^ΔuORF^ mice

| Tissue | Pathology/Histology | | wt females | $\Delta uORF$ females | wt males | $\Delta uORF$ males |
| --- | --- | --- | --- | --- | --- | --- |
| Liver | Hepatocellular vacuolation | Number<sup>1</sup><br>Grade<sup>2</sup> ± SEM<br>Significance<sup>3</sup> | 9 (9)<br>1.06 ± 0.19 | 9 (9)<br>1.06 ± 0.13 | 9 (9)<br>1.5 ± 0.20 | 9 (9)<br>0.67 ± 0.12<br>**p < 0.01** |
| | Cytoplasmic nuclear inclusions | Number<sup>1</sup><br>Grade<sup>2</sup> ± SEM<br>Significance<sup>3</sup> | 2 (9)<br>0.06 ± 0.04 | 2 (9)<br>0.14 ± 0.11<br>ns | 9 (9)<br>1.11 ± 0.14 | 5 (9)<br>0.33 ± 0.14<br>**p < 0.01** |
| | Polyploidy | Number<sup>1</sup><br>Grade<sup>2</sup> ± SEM<br>Significance<sup>3</sup> | 9 (9)<br>1.39 ± 0.11 | 9 (9)<br>1.72 ± 0.17<br>ns | 9 (9)<br>2.28 ± 0.09 | 9 (9)<br>2.06 ± 0.10<br>ns |
| | Lymphoplasmatic inflammation | Number<sup>1</sup><br>Grade<sup>2</sup> ± SEM<br>Significance<sup>3</sup> | 4 (9)<br>0.44 ± 0.19 | 8 (9)<br>1.28 ± 0.28<br>**p < 0.05** | 4 (9)<br>0.39 ± 0.22 | 7 (9)<br>0.72 ± 0.21<br>ns |
| Pancreas | Islet cell hyperplasia | Number<sup>1</sup><br>Grade<sup>2</sup> ± SEM<br>Significance<sup>3</sup> | 7 (9)<br>0.72 ± 0.17 | 4 (9)<br>0.28 ± 0.12<br>**p < 0.05** | 6 (9)<br>0.50 ± 0.17 | 1 (9)<br>0.06 ± 0.06<br>**p < 0.05** |
| | Focal acinar cell atrophy | Number<sup>1</sup><br>Grade<sup>2</sup> ± SEM<br>Significance<sup>3</sup> | 0 (9)<br>0 | 2 (9)<br>0.11 ± 0.07<br>ns | 2 (9)<br>0.17 ± 0.12 | 1 (9)<br>0.06 ± 0.06<br>ns |
| | Fatty replacement of acinar cells | Number<sup>1</sup><br>Grade<sup>2</sup> ± SEM<br>Significance<sup>3</sup> | 0 (9)<br>0 | 2 (9)<br>0.17 ± 0.12<br>ns | 2 (9)<br>0.17 ± 0.12 | 1 (9)<br>0.06 ± 0.06<br>ns |
| | Inflammation | Number<sup>1</sup><br>Grade<sup>2</sup> ± SEM<br>Significance<sup>3</sup> | 6 (9)<br>0.39 ± 0.11 | 7 (9)<br>0.61 ± 0.16<br>ns | 3 (9)<br>0.17 ± 0.08 | 6 (9)<br>0.33 ± 0.08<br>ns |
| Skeletal muscle | Regenerative muscle fibers | Number<sup>1</sup><br>No.<sup>4</sup> ± SEM<br>Significance<sup>3</sup> | 1 (8)<br>0.13 ± 0.13 | 3 (9)<br>0.78 ± 0.55<br>ns | 3 (9)<br>0.78 ± 0.40 | 6 (9)<br>6.33 ± 2.13<br>**p < 0.05** |
| | Nuclear condensation | Number<sup>1</sup><br>Grade<sup>2</sup> ± SEM<br>Significance<sup>3</sup> | 3 (9)<br>0.22 ± 0.12 | 7 (9)<br>0.78 ± 0.22<br>**p < 0.05** | 9 (9)<br>1.22 ± 0.21 | 7 (9)<br>1.22 ± 0.28 |
| | Intramuscular adipose tissue | Number<sup>1</sup><br>Grade<sup>2</sup> ± SEM<br>Significance<sup>3</sup> | 6 (9)<br>2.89 ± 1.11 | 7 (9)<br>2.24 ± 1.15<br>ns | 5 (9)<br>2.82 ± 2.93 | 6 (9)<br>8.33 ± 2.92<br>ns |
| Spleen | Lymphoid hyperplasia | Number<sup>1</sup><br>Grade<sup>2</sup> ± SEM<br>Significance<sup>3</sup> | 8 (9)<br>1.83 ± 0.26 | 9 (9)<br>1.72 ± 0.12<br>ns | 9 (9)<br>1.67 ± 0.14 | 9 (9)<br>1.72 ± 0.17<br>ns |
| | Extramedullary hematopoiesis | Number<sup>1</sup><br>Grade<sup>2</sup> ± SEM<br>Significance<sup>3</sup> | 9 (9)<br>1.50 ± 0.24 | 9 (9)<br>1.06 ± 0.16<br>ns | 9 (9)<br>1.11 ± 0.14 | 9 (9)<br>1.28 ± 0.22<br>ns |
| | Dendritic reticular cell hyperplasia | Number<sup>1</sup><br>Grade<sup>2</sup> ± SEM<br>Significance<sup>3</sup> | 4 (9)<br>0.28 ± 0.12 | 2 (9)<br>0.22 ± 0.15<br>ns | 4 (9)<br>0.22 ± 0.09 | 4 (9)<br>0.56 ± 0.28<br>ns |
| Skin | Dermal inflammation | Number<sup>1</sup><br>Grade<sup>2</sup> ± SEM<br>Significance<sup>3</sup> | 9 (9)<br>0.89 ± 0.16 | 8 (9)<br>0.44 ± 0.06<br>**p < 0.05** | 9 (9)<br>0.39 ± 0.04 | 9 (9)<br>0.39 ± 0.04 |
| Bone | Bone volume<sup>6</sup> | Bone vol. / tissue vol. (%) ± SEM | 0.56 ± 0.11 (n=5) | 0.51 ± 0.27 ns (n=5) | 3.94 ± 0.42 (n=5) | 3.69 ± 1.14 ns (n=5) |
| | Trabecular number<sup>6</sup> | Number (1/μm) ± SEM | 9.00x10<sup>-5</sup> ± 1.48x10<sup>-5</sup> (n=5) | 9.40x10<sup>-5</sup> ± 5.14x10<sup>-5</sup> ns (n=5) | 4.94 x10<sup>-4</sup> ± 5.86x10<sup>-5</sup> (n=5) | 5.66x10<sup>-4</sup> ± 1.75x10<sup>-4</sup> ns (n=5) |
| | Trabecular thickness<sup>6</sup> | Thickness (μm) ± SEM | 62.27 ± 3.97 (n=5) | 54.63 ± 10.35 ns (n=5) | 80.94 ± 4.58 (n=5) | 65.50 ± 1.64 **p < 0.05** (n=5) |
| | Trabecular separation<sup>6</sup> | Distance (μm) ± SEM | 697.55 ± 48.59 (n=5) | 777.81 ± 50.82 ns (n=5) | 469.05 ± 36.98 (n=5) | 489.07 ± 42.27 ns (n=5) |
| Blood | IGF-1 levels (plasma)<sup>9</sup> | ng/ml ± SEM<br>Significance<sup>3</sup> | 375.4 ± 17.8 (n=11) | 321.5 ± 16.4 **p < 0.05** (n=11) | 291.2 ± 21.9 (n=12) | 238.6 ± 22.5 ns (n=10) |

Mice were part of the ageing cohort and the age at analysis was 20 months for females and 22 months for males.
<sup>1</sup>Number of animals showing the pathology (out of the total number of animals analyzed).
<sup>2</sup>mean grade of the pathology as calculated from the total number of animals analyzed with 0 = absent, 1 = mild, 2 = moderate and 3 = severe.
<sup>3</sup>Statistical significance of difference found between wt and C/EBP $\beta^{\Delta uORF}$ mice from the same gender as calculated using the Student's t-test (ns = not significant).
<sup>4</sup>Mean number of regenerating muscle fibers found in five histological tissue slices per mouse. Note that a lower number is an indication for a more progressed ageing phenotype.
<sup>5</sup>mean surface area of intramuscular adipose tissue in percent of the total area of analyzed skeletal muscle tissue as calculated from the total number of animals analyzed.
<sup>6</sup>Trabecular bone parameters (percent bone volume/tissue volume; Trabecular number per mm; trabecular thickness and trabecular separation) measured by micro-CT analysis.

**Table supplement 4.** GO-term analysis of genes up-regulated in livers of old C/EBPβ^ΔuORF^ mice

| GO term | Description | p-value | FDR | Number of genes | Fold enrichment |
| --- | --- | --- | --- | --- | --- |
| GO:0009897 | External side of plasma membrane | $4.4 \times 10^{-11}$ | $5.2 \times 10^{-8}$ | 16 | 10.1 |
| GO:0042102 | Positive regulation of T cell proliferation | $2.5 \times 10^{-8}$ | $3.8 \times 10^{-5}$ | 8 | 25.4 |
| GO:0016020 | Membrane | $3.9 \times 10^{-8}$ | $4.7 \times 10^{-5}$ | 62 | 1.8 |
| GO:0031295 | T cell costimulation | $1.2 \times 10^{-7}$ | $1.8 \times 10^{-4}$ | 6 | 48.8 |
| GO:0001772 | Immunological synapse | $6.5 \times 10^{-7}$ | $7.8 \times 10^{-4}$ | 6 | 35.4 |
| GO:0006955 | Immune response | $8.1 \times 10^{-7}$ | $1.2 \times 10^{-3}$ | 11 | 8.2 |
| GO:0042113 | B cell activation | $1.1 \times 10^{-5}$ | $1.7 \times 10^{-2}$ | 5 | 35.0 |
| GO:0046641 | Positive regulation of alpha-beta T cell proliferation | $1.8 \times 10^{-5}$ | $2.7 \times 10^{-2}$ | 4 | 73.9 |
| GO:0045086 | Positive regulation of interleukine-2 biosynthetic process | $2.4 \times 10^{-5}$ | $3.5 \times 10^{-2}$ | 4 | 67.7 |

Functional annotation of genes upregulated in livers of old C/EBP $\beta^{\Delta uORF}$ female mice compared to livers of old wt female mice (FDR < 0.01; 103 from 127 genes; 24 unknown IDs) using the DAVID database (Huang et al., 2009)

**Table supplement 5.** GO-term analysis of genes down-regulated in livers of oldC/EBPβ^ΔuORF^ mice

| GO term | Description | p-value | FDR q-value | Number of genes | Fold enrichment |
| --- | --- | --- | --- | --- | --- |
| GO:0006953 | Acute phase response | $5.3 \times 10^{-6}$ | $6.2 \times 10^{-3}$ | 4 | 108.8 |
| GO:0005576 | Extracellular space | $2.5 \times 10^{-5}$ | $2.4 \times 10^{-2}$ | 9 | 6.2 |

Functional annotation of genes downregulated in livers of old C/EBP $\beta^{\Delta uORF}$ female mice compared to livers of old wt female mice (FDR < 0.01; 23 from 25 genes, 2 unknown IDs) using the DAVID database (Huang et al., 2009)

**Table supplement 6.** GO-term analysis of genes showing high inter-individual variation between livers of old wt female mice

| GO term | Description | p-value | FDR q-value | Number of genes | Fold enrichment |
| --- | --- | --- | --- | --- | --- |
| GO:0005739 | Mitochondrion | $1.2 \times 10^{-30}$ | $1.8 \times 10^{-27}$ | 236 | 2.1 |
| GO:0070062 | Extracellular exosome | $3.4 \times 10^{-29}$ | $5.1 \times 10^{-26}$ | 315 | 1.8 |
| GO:0055114 | Oxidation-reduction process | $4.4 \times 10^{-26}$ | $8.0 \times 10^{-23}$ | 124 | 2.8 |
| GO:0005743 | Mitochondrial inner membrane | $2.5 \times 10^{-24}$ | $3.7 \times 10^{-21}$ | 86 | 3.5 |
| GO:0016491 | Oxidoreductase activity | $2.6 \times 10^{-24}$ | $4.2 \times 10^{-21}$ | 113 | 2.8 |
| GO:0005783 | Endoplasmic reticulum | $1.8 \times 10^{-17}$ | $2.6 \times 10^{-14}$ | 166 | 2.0 |
| GO:0008152 | Metabolic process | $1.7 \times 10^{-12}$ | $3.1 \times 10^{-9}$ | 74 | 2.4 |
| GO:0072562 | Blood microparticle | $5.0 \times 10^{-11}$ | $7.3 \times 10^{-8}$ | 33 | 3.9 |
| GO:0006953 | Acute phase response | $1.2 \times 10^{-10}$ | $2.3 \times 10^{-7}$ | 17 | 7.4 |
| GO:0005789 | Endoplasmic reticulum membrane | $1.6 \times 10^{-10}$ | $2.4 \times 10^{-7}$ | 92 | 2.0 |
| GO:0006810 | Transport | $2.3 \times 10^{-10}$ | $4.2 \times 10^{-7}$ | 187 | 1.6 |
| GO:0043231 | Intracellular membrane-bounded organelle | $2.9 \times 10^{-10}$ | $4.3 \times 10^{-7}$ | 95 | 2.0 |
| GO:0007596 | Blood coagulation | $2.5 \times 10^{-9}$ | $4.5 \times 10^{-6}$ | 24 | 4.4 |
| GO:0042730 | Fibrinolysis | $3.0 \times 10^{-9}$ | $5.6 \times 10^{-6}$ | 11 | 11.2 |
| GO:0016020 | Membrane | $4.5 \times 10^{-9}$ | $6.6 \times 10^{-6}$ | 546 | 1.2 |
| GO:0006635 | Fatty acid beta-oxidation | $5.9 \times 10^{-8}$ | $1.1 \times 10^{-4}$ | 16 | 5.6 |
| GO:0006631 | Fatty acid metabolic process | $8.0 \times 10^{-8}$ | $1.5 \times 10^{-4}$ | 31 | 3.0 |
| GO:0031090 | Organelle membrane | $8.7 \times 10^{-8}$ | $1.3 \times 10^{-4}$ | 23 | 3.8 |
| GO:0006629 | Lipid metabolic process | $9.2 \times 10^{-8}$ | $1.7 \times 10^{-4}$ | 62 | 2.1 |
| GO:0007599 | Hemostasis | $1.2 \times 10^{-7}$ | $2.1 \times 10^{-4}$ | 16 | 5.3 |
| GO:0005777 | Peroxisome | $2.1 \times 10^{-7}$ | $3.1 \times 10^{-4}$ | 27 | 3.2 |
| GO:0006749 | Glutathione metabolic process | $3.0 \times 10^{-7}$ | $5.5 \times 10^{-4}$ | 16 | 5.0 |
| GO:0005829 | Cytosol | $4.6 \times 10^{-7}$ | $6.7 \times 10^{-4}$ | 167 | 1.5 |
| GO:0005615 | Extracellular space | $9.5 \times 10^{-7}$ | $1.4 \times 10^{-3}$ | 144 | 1.5 |
| GO:0034364 | High-density lipoprotein particle | $2.6 \times 10^{-6}$ | $3.8 \times 10^{-3}$ | 10 | 7.4 |
| GO:0005793 | Endoplasmic reticulum-Golgi intermediate compartment | $4.6 \times 10^{-6}$ | $6.7 \times 10^{-3}$ | 17 | 3.9 |
| GO:0004364 | Glutathione transferase activity | $4.8 \times 10^{-6}$ | $7.8 \times 10^{-3}$ | 12 | 5.5 |
| GO:0004129 | Cytochrome-c oxidase activity | $6.4 \times 10^{-6}$ | $1.1 \times 10^{-2}$ | 11 | 6.0 |
| GO:0003988 | Acetyl-CoA C-acetyltransferase activity | $6.9 \times 10^{-6}$ | $1.1 \times 10^{-2}$ | 6 | 15.2 |
| GO:0006099 | Tricarboxylic acid cycle | $8.7 \times 10^{-6}$ | $1.6 \times 10^{-2}$ | 11 | 5.8 |
| GO:0020037 | Heme binding | $1.1 \times 10^{-5}$ | $1.7 \times 10^{-2}$ | 29 | 2.5 |
| GO:0043022 | Ribosome binding | $1.1 \times 10^{-5}$ | $1.8 \times 10^{-2}$ | 15 | 4.1 |
| GO:0016829 | Lyase activity | $1.2 \times 10^{-5}$ | $2.0 \times 10^{-2}$ | 25 | 2.7 |
| GO:0005765 | Lysosomal membrane | $1.7 \times 10^{-5}$ | $2.5 \times 10^{-2}$ | 34 | 2.3 |
| GO:0016787 | Hydrolase activity | $2.2 \times 10^{-5}$ | $3.6 \times 10^{-2}$ | 142 | 1.4 |
| GO:0004497 | Monooxygenase activity | $2.2 \times 10^{-5}$ | $3.6 \times 10^{-2}$ | 21 | 3.0 |

Functional annotation of genes showing a high inter-individual variation between livers from old wt female mice (Coefficient of variation of transcript levels of the corresponding gene in the livers of wt mice is at least twice as big as the coefficient of variation of the transcript levels of the same gene in the livers from C/EBP $\beta^{\Delta uORF}$ mice; 1386 from 1414 genes, 28 unknown IDs) using the DAVID database (Huang et al., 2009)

**Table supplement 7.**
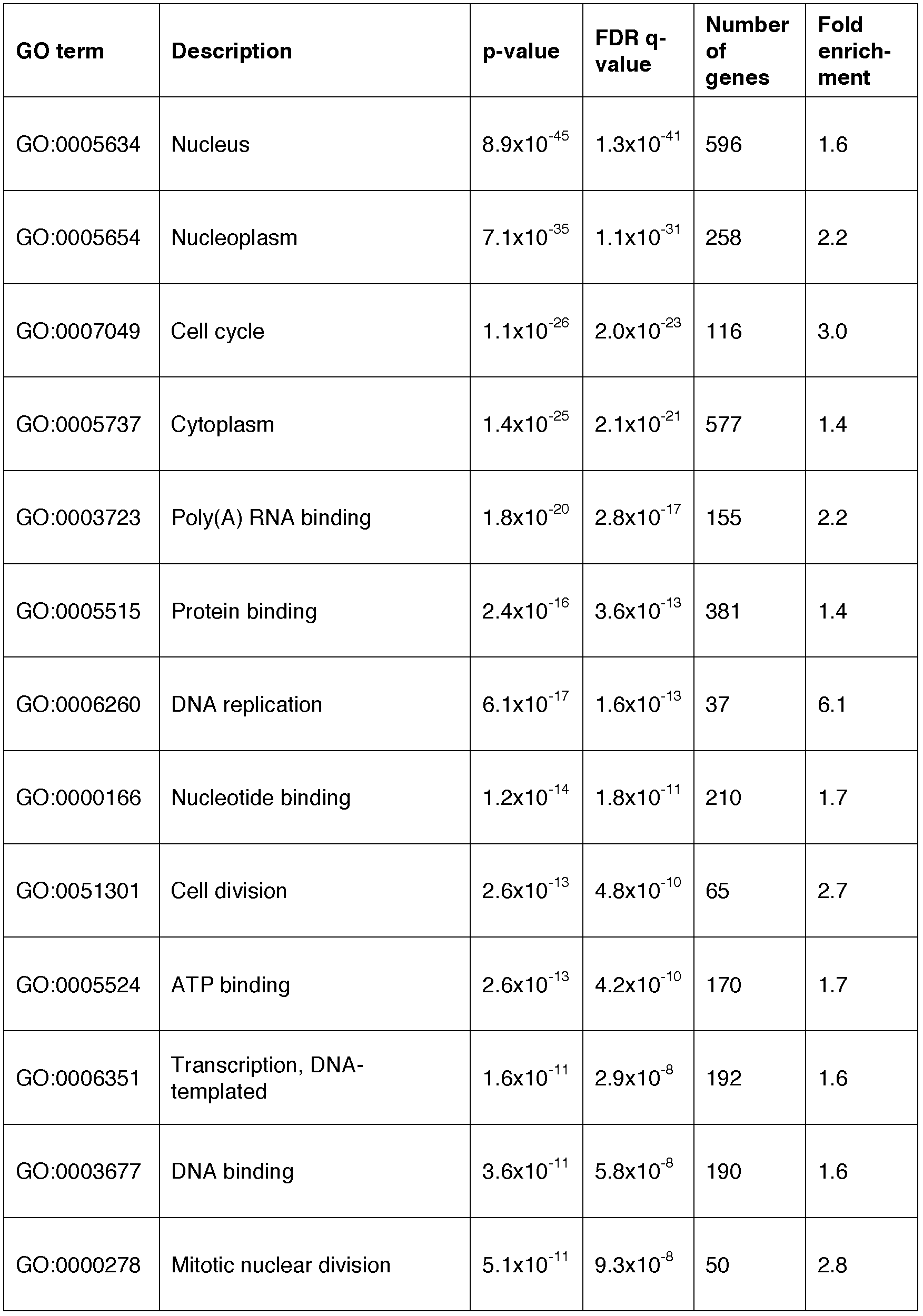
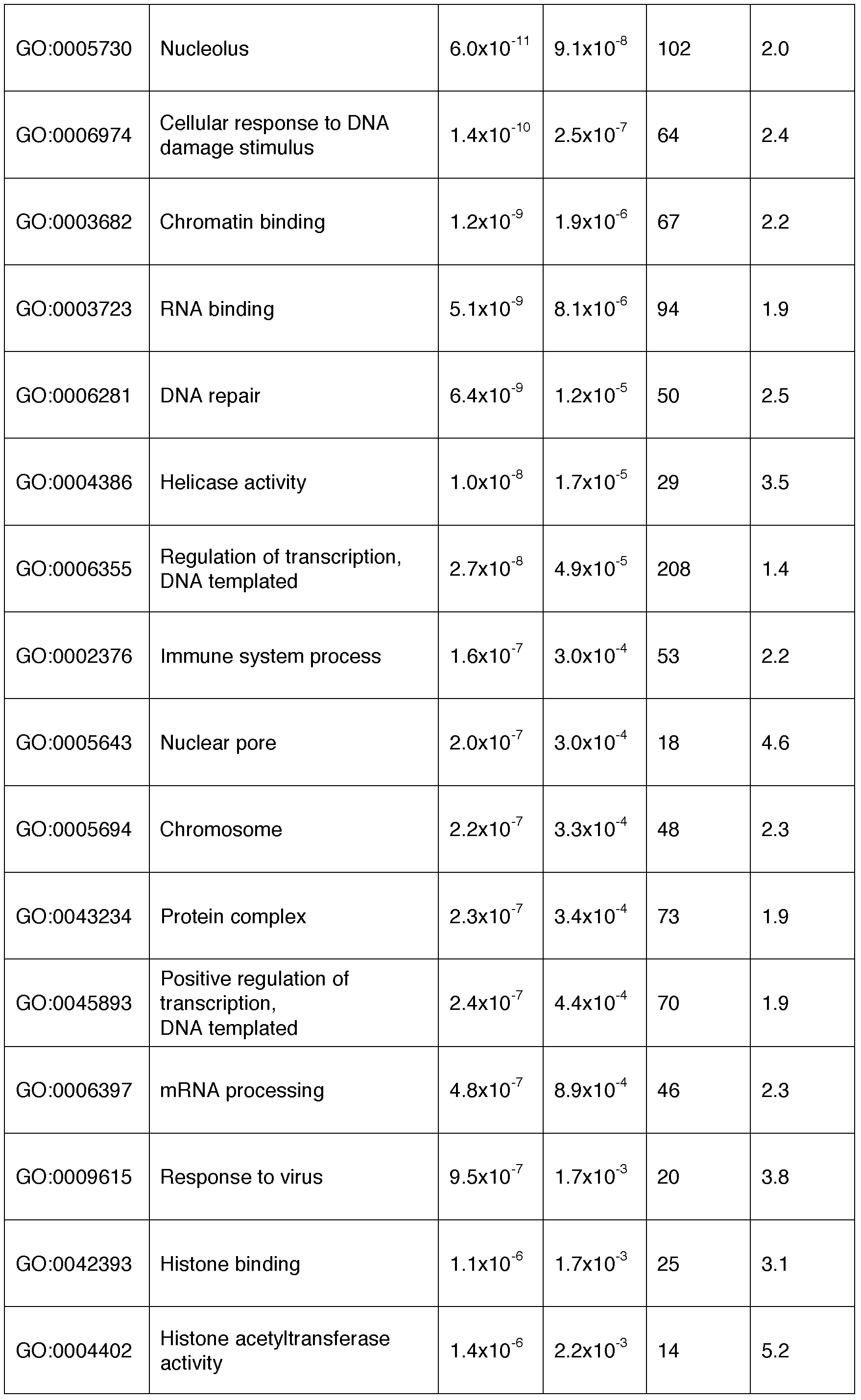
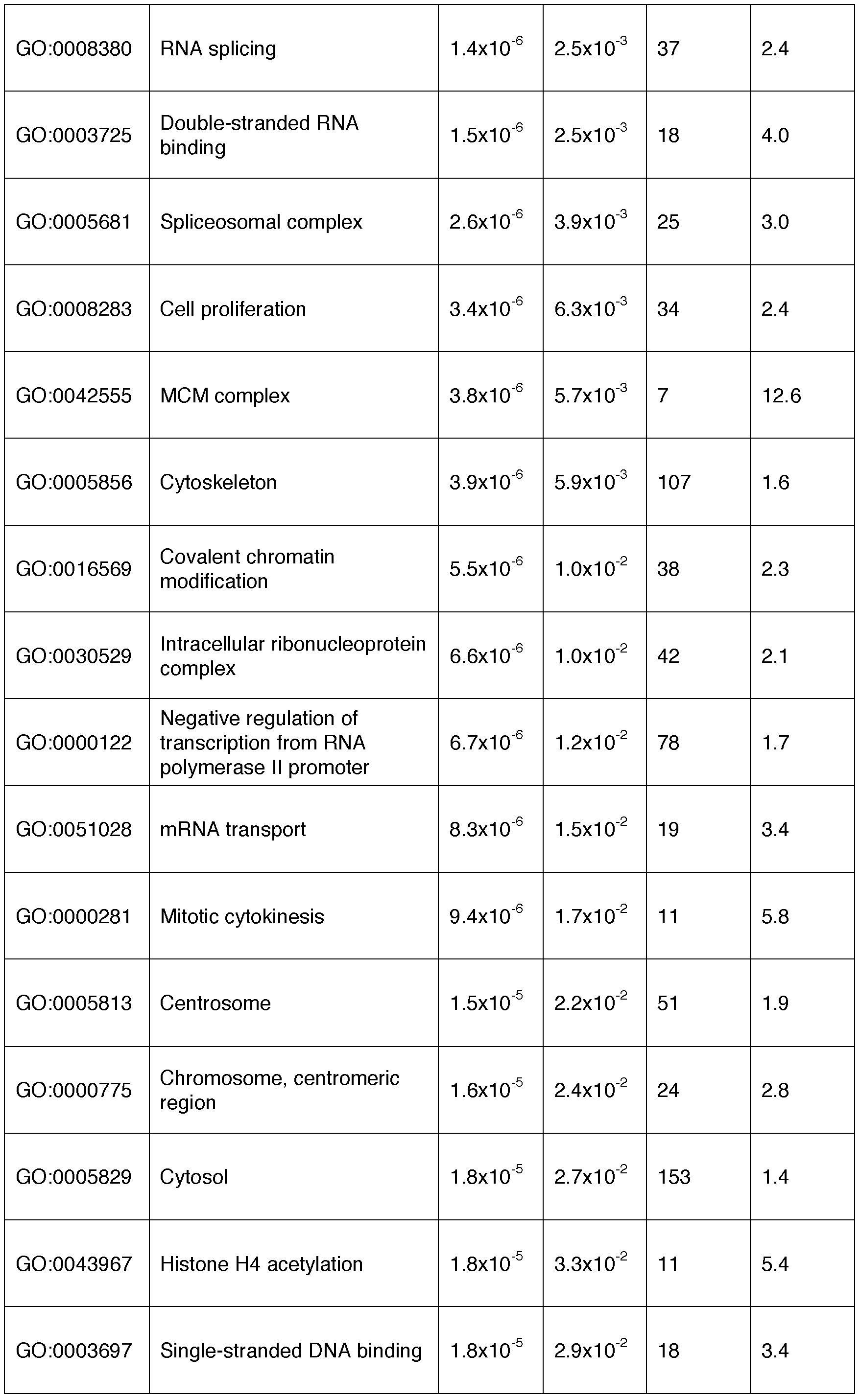
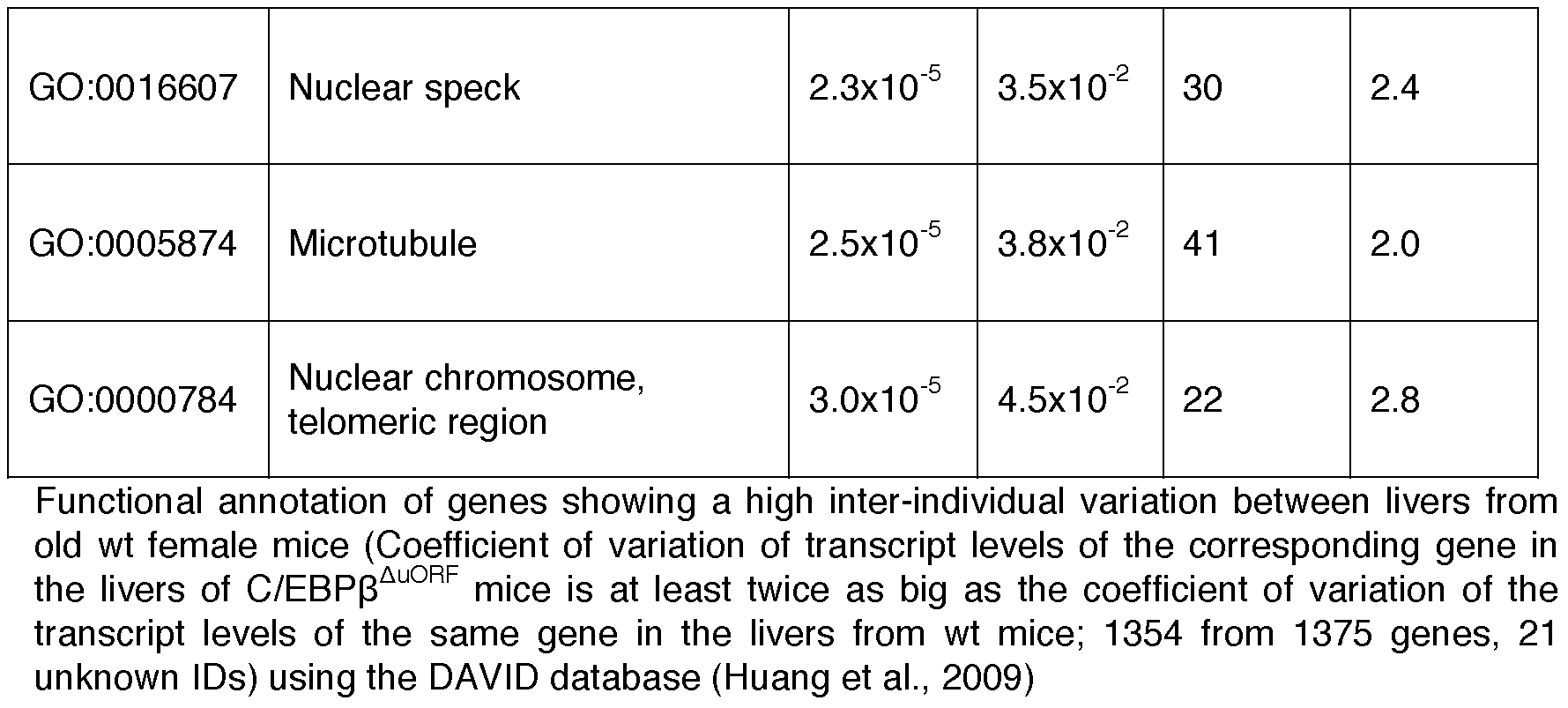
GO-term analysis of genes showing high inter-individual variation between livers of old C/EBPβ^ΔuORF^ female mice

| GO term | Description | p-value | FDR q-value | Number of genes | Fold enrichment |
| --- | --- | --- | --- | --- | --- |
| GO:0005634 | Nucleus | $8.9 \times 10^{-45}$ | $1.3 \times 10^{-41}$ | 596 | 1.6 |
| GO:0005654 | Nucleoplasm | $7.1 \times 10^{-35}$ | $1.1 \times 10^{-31}$ | 258 | 2.2 |
| GO:0007049 | Cell cycle | $1.1 \times 10^{-26}$ | $2.0 \times 10^{-23}$ | 116 | 3.0 |
| GO:0005737 | Cytoplasm | $1.4 \times 10^{-25}$ | $2.1 \times 10^{-21}$ | 577 | 1.4 |
| GO:0003723 | Poly(A) RNA binding | $1.8 \times 10^{-20}$ | $2.8 \times 10^{-17}$ | 155 | 2.2 |
| GO:0005515 | Protein binding | $2.4 \times 10^{-16}$ | $3.6 \times 10^{-13}$ | 381 | 1.4 |
| GO:0006260 | DNA replication | $6.1 \times 10^{-17}$ | $1.6 \times 10^{-13}$ | 37 | 6.1 |
| GO:0000166 | Nucleotide binding | $1.2 \times 10^{-14}$ | $1.8 \times 10^{-11}$ | 210 | 1.7 |
| GO:0051301 | Cell division | $2.6 \times 10^{-13}$ | $4.8 \times 10^{-10}$ | 65 | 2.7 |
| GO:0005524 | ATP binding | $2.6 \times 10^{-13}$ | $4.2 \times 10^{-10}$ | 170 | 1.7 |
| GO:0006351 | Transcription, DNA-templated | $1.6 \times 10^{-11}$ | $2.9 \times 10^{-8}$ | 192 | 1.6 |
| GO:0003677 | DNA binding | $3.6 \times 10^{-11}$ | $5.8 \times 10^{-8}$ | 190 | 1.6 |
| GO:0000278 | Mitotic nuclear division | $5.1 \times 10^{-11}$ | $9.3 \times 10^{-8}$ | 50 | 2.8 |
| GO:0005730 | Nucleolus | $6.0 \times 10^{-11}$ | $9.1 \times 10^{-8}$ | 102 | 2.0 |
| GO:0006974 | Cellular response to DNA damage stimulus | $1.4 \times 10^{-10}$ | $2.5 \times 10^{-7}$ | 64 | 2.4 |
| GO:0003682 | Chromatin binding | $1.2 \times 10^{-9}$ | $1.9 \times 10^{-6}$ | 67 | 2.2 |
| GO:0003723 | RNA binding | $5.1 \times 10^{-9}$ | $8.1 \times 10^{-6}$ | 94 | 1.9 |
| GO:0006281 | DNA repair | $6.4 \times 10^{-9}$ | $1.2 \times 10^{-5}$ | 50 | 2.5 |
| GO:0004386 | Helicase activity | $1.0 \times 10^{-8}$ | $1.7 \times 10^{-5}$ | 29 | 3.5 |
| GO:0006355 | Regulation of transcription, DNA templated | $2.7 \times 10^{-8}$ | $4.9 \times 10^{-5}$ | 208 | 1.4 |
| GO:0002376 | Immune system process | $1.6 \times 10^{-7}$ | $3.0 \times 10^{-4}$ | 53 | 2.2 |
| GO:0005643 | Nuclear pore | $2.0 \times 10^{-7}$ | $3.0 \times 10^{-4}$ | 18 | 4.6 |
| GO:0005694 | Chromosome | $2.2 \times 10^{-7}$ | $3.3 \times 10^{-4}$ | 48 | 2.3 |
| GO:0043234 | Protein complex | $2.3 \times 10^{-7}$ | $3.4 \times 10^{-4}$ | 73 | 1.9 |
| GO:0045893 | Positive regulation of transcription, DNA templated | $2.4 \times 10^{-7}$ | $4.4 \times 10^{-4}$ | 70 | 1.9 |
| GO:0006397 | mRNA processing | $4.8 \times 10^{-7}$ | $8.9 \times 10^{-4}$ | 46 | 2.3 |
| GO:0009615 | Response to virus | $9.5 \times 10^{-7}$ | $1.7 \times 10^{-3}$ | 20 | 3.8 |
| GO:0042393 | Histone binding | $1.1 \times 10^{-6}$ | $1.7 \times 10^{-3}$ | 25 | 3.1 |
| GO:0004402 | Histone acetyltransferase activity | $1.4 \times 10^{-6}$ | $2.2 \times 10^{-3}$ | 14 | 5.2 |
| GO:0008380 | RNA splicing | $1.4 \times 10^{-6}$ | $2.5 \times 10^{-3}$ | 37 | 2.4 |
| GO:0003725 | Double-stranded RNA binding | $1.5 \times 10^{-6}$ | $2.5 \times 10^{-3}$ | 18 | 4.0 |
| GO:0005681 | Spliceosomal complex | $2.6 \times 10^{-6}$ | $3.9 \times 10^{-3}$ | 25 | 3.0 |
| GO:0008283 | Cell proliferation | $3.4 \times 10^{-6}$ | $6.3 \times 10^{-3}$ | 34 | 2.4 |
| GO:0042555 | MCM complex | $3.8 \times 10^{-6}$ | $5.7 \times 10^{-3}$ | 7 | 12.6 |
| GO:0005856 | Cytoskeleton | $3.9 \times 10^{-6}$ | $5.9 \times 10^{-3}$ | 107 | 1.6 |
| GO:0016569 | Covalent chromatin modification | $5.5 \times 10^{-6}$ | $1.0 \times 10^{-2}$ | 38 | 2.3 |
| GO:0030529 | Intracellular ribonucleoprotein complex | $6.6 \times 10^{-6}$ | $1.0 \times 10^{-2}$ | 42 | 2.1 |
| GO:0000122 | Negative regulation of transcription from RNA polymerase II promoter | $6.7 \times 10^{-6}$ | $1.2 \times 10^{-2}$ | 78 | 1.7 |
| GO:0051028 | mRNA transport | $8.3 \times 10^{-6}$ | $1.5 \times 10^{-2}$ | 19 | 3.4 |
| GO:0000281 | Mitotic cytokinesis | $9.4 \times 10^{-6}$ | $1.7 \times 10^{-2}$ | 11 | 5.8 |
| GO:0005813 | Centrosome | $1.5 \times 10^{-5}$ | $2.2 \times 10^{-2}$ | 51 | 1.9 |
| GO:0000775 | Chromosome, centromeric region | $1.6 \times 10^{-5}$ | $2.4 \times 10^{-2}$ | 24 | 2.8 |
| GO:0005829 | Cytosol | $1.8 \times 10^{-5}$ | $2.7 \times 10^{-2}$ | 153 | 1.4 |
| GO:0043967 | Histone H4 acetylation | $1.8 \times 10^{-5}$ | $3.3 \times 10^{-2}$ | 11 | 5.4 |
| GO:0003697 | Single-stranded DNA binding | $1.8 \times 10^{-5}$ | $2.9 \times 10^{-2}$ | 18 | 3.4 |
| GO:0016607 | Nuclear speck | $2.3 \times 10^{-5}$ | $3.5 \times 10^{-2}$ | 30 | 2.4 |
| GO:0005874 | Microtubule | $2.5 \times 10^{-5}$ | $3.8 \times 10^{-2}$ | 41 | 2.0 |
| GO:0000784 | Nuclear chromosome,<br>telomeric region | $3.0 \times 10^{-5}$ | $4.5 \times 10^{-2}$ | 22 | 2.8 |

Functional annotation of genes showing a high inter-individual variation between livers from old wt female mice (Coefficient of variation of transcript levels of the corresponding gene in the livers of C/EBP $\beta^{\Delta uORF}$ mice is at least twice as big as the coefficient of variation of the transcript levels of the same gene in the livers from wt mice; 1354 from 1375 genes, 21 unknown IDs) using the DAVID database (Huang et al., 2009)

## References

Albert, V., & Hall, M. N. (2015). Reduced C/EBPbeta-LIP translation improves metabolic health. EMBO Rep, 16(8), 881–882. doi:10.15252/embr.201540757

Anand, S., Ebner, J., Warren, C. B., Raam, M. S., Piliang, M., Billings, S. D., & Maytin, E. V. (2014). C/EBP transcription factors in human squamous cell carcinoma: selective changes in expression of isoforms correlate with the neoplastic state. PLoS One, 9(11), e112073. doi:10.1371/journal.pone.0112073

Anisimov, V. N., Zabezhinski, M. A., Popovich, I. G., Piskunova, T. S., Semenchenko, A. V., Tyndyk, M. L., … Blagosklonny, M. V. (2011). Rapamycin increases lifespan and inhibits spontaneous tumorigenesis in inbred female mice. Cell Cycle, 10(24), 4230–4236. doi:10.4161/cc.10.24.18486

Arnal-Estape, A., Tarragona, M., Morales, M., Guiu, M., Nadal, C., Massague, J., & Gomis, R. R. (2010). HER2 silences tumor suppression in breast cancer cells by switching expression of C/EBPss isoforms. Cancer Res, 70(23), 9927–9936. doi:10.1158/0008–5472.CAN-10–0869

Augustine, J. J., Bodziak, K. A., & Hricik, D. E. (2007). Use of sirolimus in solid organ transplantation. Drugs, 67(3), 369–391.

Barreto, G., Huang, T. T., & Giffard, R. G. (2010). Age-related defects in sensorimotor activity, spatial learning, and memory in C57BL/6 mice. J Neurosurg Anesthesiol, 22(3), 214–219. doi:10.1097/ANA.0b013e3181d56c98

Barzilai, N., Banerjee, S., Hawkins, M., Chen, W., & Rossetti, L. (1998). Caloric restriction reverses hepatic insulin resistance in aging rats by decreasing visceral fat. J Clin Invest, 101(7), 1353–1361. doi:10.1172/JCI485

Begay, V., Smink, J. J., Loddenkemper, C., Zimmermann, K., Rudolph, C., Scheller, M., … Leutz, A. (2015). Deregulation of the endogenous C/EBPbeta LIP isoform predisposes to tumorigenesis. J Mol Med (Berl), 93(1), 39–49. doi:10.1007/s00109–014-1215–5

Bitto, A., Ito, T. K., Pineda, V. V., LeTexier, N. J., Huang, H. Z., Sutlief, E., … Kaeberlein, M. (2016). Transient rapamycin treatment can increase lifespan and healthspan in middle-aged mice. Elife, 5. doi:10.7554/eLife.16351

Breese, C. R., Ingram, R. L., & Sonntag, W. E. (1991). Influence of age and long-term dietary restriction on plasma insulin-like growth factor-1 (IGF-1), IGF-1 gene expression, and IGF-1 binding proteins. J Gerontol, 46(5), B180–187.

Brooks, S. P., & Dunnett, S. B. (2009). Tests to assess motor phenotype in mice: a user’s guide. Nat Rev Neurosci, 10(7), 519–529. doi:10.1038/nrn2652

Brown-Borg, H. M. (2009). Hormonal control of aging in rodents: the somatotropic axis. Mol Cell Endocrinol, 299(1), 64–71. doi:S0303-7207(08)00285–2 [pii] 10.1016/j.mce.2008.07.001

Brown-Borg, H. M., Borg, K. E., Meliska, C. J., & Bartke, A. (1996). Dwarf mice and the ageing process. Nature, 384(6604), 33. doi:10.1038/384033a0

Calkhoven, C. F., Muller, C., & Leutz, A. (2000). Translational control of C/EBPalpha and C/EBPbeta isoform expression. Genes Dev, 14(15), 1920–1932.

Colman, R. J., Anderson, R. M., Johnson, S. C., Kastman, E. K., Kosmatka, K. J., Beasley, T. M., … Weindruch, R. (2009). Caloric restriction delays disease onset and mortality in rhesus monkeys. Science, 325(5937), 201–204. doi:10.1126/science.1173635

de Jong, T. (2018). Processed data and R script for analysis and visualization of transcriptome variability. Retrieved from http://www.genomes.nl/CEBPB_delta_uORF/

de Oliveira, M. A., Martins, E. M. F., Wang, Q., Sonis, S., Demetri, G., George, S., … Treister, N. S. (2011). Clinical presentation and management of mTOR inhibitor-associated stomatitis. Oral Oncol, 47(10), 998–1003. doi:10.1016/j.oraloncology.2011.08.009

Demontis, F., Piccirillo, R., Goldberg, A. L., & Perrimon, N. (2013). Mechanisms of skeletal muscle aging: insights from Drosophila and mammalian models. Dis Model Mech, 6(6), 1339–1352. doi:10.1242/dmm.012559

Descombes, P., & Schibler, U. (1991). A liver-enriched transcriptional activator protein, LAP, and a transcriptional inhibitory protein, LIP, are translated from the same mRNA. Cell, 67(3), 569–579. doi:0092-8674(91)90531–3 [pii]

Desvergne, B., Michalik, L., & Wahli, W. (2006). Transcriptional regulation of metabolism. Physiol Rev, 86(2), 465–514. doi:10.1152/physrev.00025.2005

Dobin, A., Davis, C. A., Schlesinger, F., Drenkow, J., Zaleski, C., Jha, S., … Gingeras, T. R. (2013). STAR: ultrafast universal RNA-seq aligner. Bioinformatics, 29(1), 15–21. doi:10.1093/bioinformatics/bts635

Fang, Y., Westbrook, R., Hill, C., Boparai, R. K., Arum, O., Spong, A., … Bartke, A. (2013). Duration of rapamycin treatment has differential effects on metabolism in mice. Cell Metab, 17(3), 456–462. doi:10.1016/j.cmet.2013.02.008

Fok, W. C., Chen, Y., Bokov, A., Zhang, Y., Salmon, A. B., Diaz, V., … Richardson, A. (2014). Mice fed rapamycin have an increase in lifespan associated with major changes in the liver transcriptome. PLoS One, 9(1), e83988. doi:10.1371/journal.pone.0083988

Haas, S. C., Huber, R., Gutsch, R., Kandemir, J. D., Cappello, C., Krauter, J., … Brand, K. (2010). ITD‐ and FL-induced FLT3 signal transduction leads to increased C/EBPbeta-LIP expression and LIP/LAP ratio by different signalling modules. Br J Haematol, 148(5), 777–790. doi:10.1111/j.1365–2141.2009.08012.x

Hakim, F. T., Flomerfelt, F. A., Boyiadzis, M., & Gress, R. E. (2004). Aging, immunity and cancer. Curr Opin Immunol, 16(2), 151–156. doi:10.1016/j.coi.2004.01.009

Hampton, A. L., Hish, G. A., Aslam, M. N., Rothman, E. D., Bergin, I. L., Patterson, K. A., … Rush, H. G. (2012). Progression of ulcerative dermatitis lesions in C57BL/6Crl mice and the development of a scoring system for dermatitis lesions. J Am Assoc Lab Anim Sci, 51(5), 586–593.

Harrison, D. E., Strong, R., Sharp, Z. D., Nelson, J. F., Astle, C. M., Flurkey, K., … Miller, R. A. (2009). Rapamycin fed late in life extends lifespan in genetically heterogeneous mice. Nature, 460(7253), 392–395. doi:10.1038/nature08221

Hsieh, C. C., Xiong, W., Xie, Q., Rabek, J. P., Scott, S. G., An, M. R., … Papaconstantinou, J. (1998). Effects of age on the posttranscriptional regulation of CCAAT/enhancer binding protein alpha and CCAAT/enhancer binding protein beta isoform synthesis in control and LPS-treated livers. Mol Biol Cell, 9(6), 1479–1494.

Huang da, W., Sherman, B. T., & Lempicki, R. A. (2009). Systematic and integrative analysis of large gene lists using DAVID bioinformatics resources. Nat Protoc, 4(1), 44–57. doi:10.1038/nprot.2008.211

Johnson, S. C., Rabinovitch, P. S., & Kaeberlein, M. (2013). mTOR is a key modulator of ageing and age-related disease. Nature, 493(7432), 338–345. doi:10.1038/nature11861

Jundt, F., Raetzel, N., Muller, C., Calkhoven, C. F., Kley, K., Mathas, S., … Dorken, B. (2005). A rapamycin derivative (everolimus) controls proliferation through down-regulation of truncated CCAAT enhancer binding protein {beta} and NF-{kappa}B activity in Hodgkin and anaplastic large cell lymphomas. Blood, 106(5), 1801–1807. doi:2004–11-4513 [pii] 10.1182/blood-2004–11-4513

Kaeberlein, M., Rabinovitch, P. S., & Martin, G. M. (2015). Healthy aging: The ultimate preventative medicine. Science, 350(6265), 1191–1193. doi:10.1126/science.aad3267

Kang, T. W., Yevsa, T., Woller, N., Hoenicke, L., Wuestefeld, T., Dauch, D., … Zender, L. (2011). Senescence surveillance of pre-malignant hepatocytes limits liver cancer development. Nature, 479(7374), 547–551. doi:10.1038/nature10599

Karagiannides, I., Tchkonia, T., Dobson, D. E., Steppan, C. M., Cummins, P., Chan, G., … Kirkland, J. L. (2001). Altered expression of C/EBP family members results in decreased adipogenesis with aging. Am J Physiol Regul Integr Comp Physiol, 280(6), R1772–1780.

Komarova, E. A., Antoch, M. P., Novototskaya, L. R., Chernova, O. B., Paszkiewicz, G., Leontieva, O. V., … Gudkov, A. V. (2012). Rapamycin extends lifespan and delays tumorigenesis in heterozygous p53+/- mice. Aging (Albany NY), 4(10), 709–714. doi:10.18632/aging.100498

Lamming, D. W., Ye, L., Katajisto, P., Goncalves, M. D., Saitoh, M., Stevens, D. M., … Baur, J. A. (2012). Rapamycin-induced insulin resistance is mediated by mTORC2 loss and uncoupled from longevity. Science, 335(6076), 1638–1643. doi:10.1126/science.1215135

Lee, J. S., Ward, W. O., Ren, H., Vallanat, B., Darlington, G. J., Han, E. S., … Corton, J. C. (2012). Meta-analysis of gene expression in the mouse liver reveals biomarkers associated with inflammation increased early during aging. Mech Ageing Dev, 133(7), 467–478. doi:10.1016/j.mad.2012.05.006

Liu, J., Ibi, D., Taniguchi, K., Lee, J., Herrema, H., Akosman, B., … Ozcan, U. (2016). Inflammation Improves Glucose Homeostasis through IKKbeta-XBP1s Interaction. Cell, 167(4), 1052–1066 e1018. doi:10.1016/j.cell.2016.10.015

Martin-Montalvo, A., Mercken, E. M., Mitchell, S. J., Palacios, H. H., Mote, P. L., Scheibye-Knudsen, M., … de Cabo, R. (2013). Metformin improves healthspan and lifespan in mice. Nat Commun, 4, 2192. doi:10.1038/ncomms3192

Mattison, J. A., Roth, G. S., Beasley, T. M., Tilmont, E. M., Handy, A. M., Herbert, R. L., … de Cabo, R. (2012). Impact of caloric restriction on health and survival in rhesus monkeys from the NIA study. Nature, 489(7415), 318–321. doi:10.1038/nature11432

McCarthy, S. D., Roche, J. F., & Forde, N. (2012). Temporal changes in endometrial gene expression and protein localization of members of the IGF family in cattle: effects of progesterone and pregnancy. Physiol Genomics, 44(2), 130–140. doi:10.1152/physiolgenomics.00106.2011

Miller, R. A., Harrison, D. E., Astle, C. M., Baur, J. A., Boyd, A. R., de Cabo, R., … Strong, R. (2011). Rapamycin, but not resveratrol or simvastatin, extends life span of genetically heterogeneous mice. J Gerontol A Biol Sci Med Sci, 66(2), 191–201. doi:10.1093/gerona/glq178

Mitchell, S. J., Madrigal-Matute, J., Scheibye-Knudsen, M., Fang, E., Aon, M., Gonzalez-Reyes, J. A., … de Cabo, R. (2016). Effects of Sex, Strain, and Energy Intake on Hallmarks of Aging in Mice. Cell Metab, 23(6), 1093–1112. doi:10.1016/j.cmet.2016.05.027

Müller, C., de Jong, T, Guryev, V, Calkhoven, CF. (2018). Transcriptome profiling of liver samples of C/EBPβΔuORF mice. Retrieved from: https://www.ebi.ac.uk/arrayexpress/

Neff, F., Flores-Dominguez, D., Ryan, D. P., Horsch, M., Schroder, S., Adler, T., … Ehninger, D. (2013). Rapamycin extends murine lifespan but has limited effects on aging. J Clin Invest, 123(8), 3272–3291. doi:10.1172/JCI67674

Park, B. H., Kook, S., Lee, S., Jeong, J. H., Brufsky, A., & Lee, B. C. (2013). An isoform of C/EBPbeta, LIP, regulates expression of the chemokine receptor CXCR4 and modulates breast cancer cell migration. J Biol Chem, 288(40), 28656–28667. doi:10.1074/jbc.M113.509505

Raught, B., Gingras, A. C., James, A., Medina, D., Sonenberg, N., & Rosen, J. M. (1996). Expression of a translationally regulated, dominant-negative CCAAT/enhancer-binding protein beta isoform and up-regulation of the eukaryotic translation initiation factor 2alpha are correlated with neoplastic transformation of mammary epithelial cells. Cancer Res, 56(19), 4382–4386.

Reimand, J., Arak, T., Adler, P., Kolberg, L., Reisberg, S., Peterson, H., & Vilo, J. (2016). g:Profiler-a web server for functional interpretation of gene lists (2016 update). Nucleic Acids Res, 44(W1), W83–89. doi:10.1093/nar/gkw199

Robinson, M. D., McCarthy, D. J., & Smyth, G. K. (2010). edgeR: a Bioconductor package for differential expression analysis of digital gene expression data. Bioinformatics, 26(1), 139–140. doi:10.1093/bioinformatics/btp616

Roesler, W. J. (2001). The role of C/EBP in nutrient and hormonal regulation of gene expression. Annu Rev Nutr, 21, 141–165. doi:10.1146/annurev.nutr.21.1.141 [pii]

Selman, C., Tullet, J. M., Wieser, D., Irvine, E., Lingard, S. J., Choudhury, A. I., … Withers, D. J. (2009). Ribosomal protein S6 kinase 1 signaling regulates mammalian life span. Science, 326(5949), 140–144. doi:326/5949/140 [pii] 10.1126/science.1177221

Serrano, M. (2016). Unraveling the links between cancer and aging. Carcinogenesis, 37(2), 107. doi:10.1093/carcin/bgv100

Singh, P., Coskun, Z. Z., Goode, C., Dean, A., Thompson-Snipes, L., & Darlington, G. (2008). Lymphoid neogenesis and immune infiltration in aged liver. Hepatology, 47(5), 1680–1690. doi:10.1002/hep.22224

Timchenko, L. T., Salisbury, E., Wang, G. L., Nguyen, H., Albrecht, J. H., Hershey, J. W., & Timchenko, N. A. (2006). Age-specific CUGBP1-eIF2 complex increases translation of CCAAT/enhancer-binding protein beta in old liver. J Biol Chem, 281(43), 32806–32819. doi:10.1074/jbc.M605701200

Weindruch, R., & Walford, R. L. (1982). Dietary restriction in mice beginning at 1 year of age: effect on life-span and spontaneous cancer incidence. Science, 215(4538), 1415–1418.

Wethmar, K., Begay, V., Smink, J. J., Zaragoza, K., Wiesenthal, V., Dorken, B., … Leutz, A. (2010). C/EBPbetaDeltauORF mice‐‐a genetic model for uORF-mediated translational control in mammals. Genes Dev, 24(1), 15–20. doi:24/1/15 [pii] 10.1101/gad.557910

White, R. R., Milholland, B., MacRae, S. L., Lin, M., Zheng, D., & Vijg, J. (2015). Comprehensive transcriptional landscape of aging mouse liver. BMC Genomics, 16, 899. doi:10.1186/s12864–015-2061–8

Wilkinson, J. E., Burmeister, L., Brooks, S. V., Chan, C. C., Friedline, S., Harrison, D. E., … Miller, R. A. (2012). Rapamycin slows aging in mice. Aging Cell, 11(4), 675–682. doi:10.1111/j.1474–9726.2012.00832.x

Wu, J. J., Liu, J., Chen, E. B., Wang, J. J., Cao, L., Narayan, N., … Finkel, T. (2013). Increased Mammalian Lifespan and a Segmental and Tissue-Specific Slowing of Aging after Genetic Reduction of mTOR Expression. Cell Rep, 4(5), 913–920. doi:10.1016/j.celrep.2013.07.030

Zahnow, C. A., Younes, P., Laucirica, R., & Rosen, J. M. (1997). Overexpression of C/EBPbeta-LIP, a naturally occurring, dominant-negative transcription factor, in human breast cancer. J Natl Cancer Inst, 89(24), 1887–1891.

Zaini, M. A., Muller, C., Ackermann, T., Reinshagen, J., Kortman, G., Pless, O., & Calkhoven, C. F. (2017). A screening strategy for the discovery of drugs that reduce C/EBPbeta-LIP translation with potential calorie restriction mimetic properties. Sci Rep, 7, 42603. doi:10.1038/srep42603

Zhang, H. M., Diaz, V., Walsh, M. E., & Zhang, Y. (2017). Moderate lifelong overexpression of tuberous sclerosis complex 1 (TSC1) improves health and survival in mice. Sci Rep, 7(1), 834. doi:10.1038/s41598–017-00970–7

Zhang, Y., Bokov, A., Gelfond, J., Soto, V., Ikeno, Y., Hubbard, G., … Fischer, K. (2014). Rapamycin extends life and health in C57BL/6 mice. J Gerontol A Biol Sci Med Sci, 69(2), 119–130. doi:10.1093/gerona/glt056

Zidek, L. M., Ackermann, T., Hartleben, G., Eichwald, S., Kortman, G., Kiehntopf, M., … Calkhoven, C. F. (2015). Deficiency in mTORC1-controlled C/EBPbeta-mRNA translation improves metabolic health in mice. EMBO Rep, 16(8), 1022–1036. doi:10.15252/embr.201439837

